# COMMD10 Regulates Endosomal Recycling of Epithelial Sodium Channel (ENaC)

**DOI:** 10.1101/2024.06.11.598390

**Authors:** Sahib R. Rasulov, Fiona J. McDonald

## Abstract

The epithelial sodium channel (ENaC) plays an essential role in the regulation of sodium transport in distal nephron. ENaC cell surface population of renal principal cells is tightly regulated by hormones such as aldosterone and vasopressin through protein trafficking pathways that translocate ENaC to and from the cell surface. Internalized ENaC from the plasma membrane follows the degradative pathway promoted by ubiquitin fusion or the recycling pathway after deubiquitination and sorting on early endosomes. The mechanism by which ENaC is recycled back to the plasma membrane through regulated recycling is less known. Here, we show that regulated recycling of ENaC is strictly dependent on COMMD10 and localization pattern of COMMD10 on Rab5, −7, and −11 vesicles is similar to that of WASH and Arp2/3. Furthermore, here we report that COMMD1 and −10 protein levels are regulated by aldosterone and calcium. This study proposes that for regulated recycling of ENaC, conventional endosomal sorting and recycling complexes such as CCC complex are recruited.

**Highlights:** COMMD10 alters ENaC cell surface population through the regulated recycling pathway. COMMD10 localizes on Rab5-, Rab7-, and Rab11-positive endosomes in a similar pattern as endosomal actin polymerization complexes WASH and Arp2/3. Aldosterone downregulates COMMD1 and −10 protein levels while calcium upregulates COMMD10 protein level.

**Graphical Abstract:** 

## Introduction

ENaC transports Na^+^ across tight epithelia including the distal nephron (Garty & Palmer, 1997; Hamm *et al*., 2010) and therefore, controlling ENaC at the cell surface is a major determinant of extracellular fluid volume and consequently blood pressure (Pratt, 2005; Rotin & Schild, 2008). The surface density of active ENaC is regulated by proteases through channel open probability, aldosterone through transcriptional pathway, and trafficking through endocytosis and recycling pathways (Girardet *et al*., 1986; Debonneville *et al*., 2001; Loffing *et al*., 2001; Abriel & Staub, 2005; Butterworth *et al*., 2005).

According to a recent model, three recycling pathways from endosomes to the plasma membrane (PM) have been classified: constitutive, regulated, and slow recycling (Cullen & Steinberg, 2018; Simonetti & Cullen, 2019). The constitutive and regulated recycling comprise the fast recycling pathway (utilizing Rab4-positive vesicles), while the recycling through the endosomal recycling compartment (ERC) is known as the slow recycling pathway (utilizing Rab11-positive vesicles) (Rea & James, 1997; Pessin *et al*., 1999; Lampson *et al*., 2001). A number of membrane proteins such as ion channels and signaling receptors escape the degradation pathways after endocytosis, and are sorted into recycling tubules on Rab5- and Rab7-positive endosomes (Cao *et al*., 1999; Hanyaloglu *et al*., 2005; Hanyaloglu & Zastrow, 2008; Millman *et al*., 2008; Yudowski *et al*., 2009) that are regulated in a cAMP/PKA-dependent manner (Vistein & Puthenveedu, 2013; Bowman *et al*., 2016). The regulated recycling pathway requires the interaction of multiple endosomal sorting complexes such as WASH, Arp2/3, CCC, retromer, and retriever (Puthenveedu *et al*., 2010; Temkin *et al*., 2011; McNally *et al*., 2017; Singla *et al*., 2019). According to a model describing the role of the CCC complex (COMMD1-10/CCDC22/CCDC93) in endosomal sorting, the early endosome marker Rab5 recruits VPS34, the class III PI3-K, onto early endosomes to generate endosomal PI(3)P (Christoforidis *et al*., 1999) which creates a base for WASH complex recruitment (Singla *et al*., 2019). WASH complex, an activator of Arp2/3, promotes endosomal F-actin formation to stabilize tubules for cargo entry (Linardopoulou *et al*., 2007), but also recruits the CCC complex (Phillips-Krawczak *et al*., 2015). After its recruitment, the CCC complex functions as a negative regulator of the WASH complex, disassembling the WASH complex from the endosomal domain (Singla *et al*., 2019). Probably, disassembly of WASH complex is required for the prevention of further endosomal F-actin formation and thereby for the release of actin-stabilized tubules. Furthermore, calcium-regulated gene COMMD5 was shown to hook endosomes to cytoskeleton and thereby to play another role in translocation of recycling vesicles to the PM (Campion *et al*., 2018).

CCC complex stabilization is highly dependent on its subunits. Thus, silencing any of the CCC complex subunits, including COMMD proteins (*<u>CO</u>*pper *<u>M</u>*etabolism *<u>M</u>*URR1 or COMM domain containing), results in CCC complex destabilization and this leads to more WASH complex accumulation on endosomes which generates more endosomal F-actin on the sorting domains and this results in longer stabilization of recycling tubules and thereby trapping of cargoes on endosomes (Singla *et al*., 2019). However, at what step the CCC complex is recruited onto endosomes to negatively regulate the WASH complex is a fundamental cell biological question that is still unanswered. Conversely, increased CCC complex activity is expected to impair WASH/Arp2/3 function reducing endosomal F-actin polymerization and thereby reducing tubule stabilization. This was confirmed with a report demonstrating that like COMMD1 knockdown (KD), COMMD1 overexpression also resulted in an accumulation of ATP7B on early endosomes (Stewart *et al*., 2019) which is consistent with our previous reports demonstrating overexpressed COMMD1, −3 and −9 downregulate ENaC cell surface activity (Ke *et al*., 2010; Liu *et al*., 2013). We have also previously reported that all COMMD1-10 family proteins interact with ENaC (Ke *et al*., 2010; Liu *et al*., 2013) and the knockdown of COMMD10 also reduces ENaC *Isc* (Ware *et al*., 2018). We also observed that both the overexpression of a subunit of WASH complex - WASHC1 and knockdown of another subunit of WASH complex, Fam21 (or WASHC2) reduces ENaC *Isc* (McDonald lab, unpublished work) suggesting that the WASH complex is required in a steady-state level for normal stabilization of ENaC-loaded tubules. Further, we have reported that COMMD10 KD reduces ENaC cell surface activity and this is mediated by reducing ENaC cell surface population (Ware *et al*., 2018). But the pathway by which COMMD10 reduces ENaC cell surface population is unknown.

In this study, we show that COMMD10 alters ENaC cell surface population through the regulated recycling pathway. We also demonstrate that COMMD10 localizes on Rab5-, Rab7-, and Rab11-positive endosomes in a similar pattern as endosomal actin polymerization complexes WASH and Arp2/3 supporting a role for COMMD10 in endosomal sorting of ENaC as a part of the CCC complex. Furthermore, here we report that COMMD1 and −10 protein levels are regulated by aldosterone and calcium.

## Results

### 1. COMMD10 knockdown impairs regulated recycling of ENaC

It was shown that silencing a subunit of the CCC complex destabilizes the CCC complex and thereby its other subunits (Phillips-Krawczak et al., 2015; Fedoseienko et al., 2018) as this was observed with reduced COMMD1 level in COMMD10 KD mCCDcl1 cells (Figure S1A,B) consistent with our previous report showing that COMMD10 KD reduces COMMD1 protein levels in FRT cells (McDonald lab, unpublished work). COMMD10 KD does not increase endocytosis rate but it results in intracellular ENaC accumulation after endocytosis (Figure S1C-F) revealing the reason of the reduction in ENaC population at the surface of COMMD10 KD cells reported in (Ware *et al*., 2018). To assess the effect of silencing of COMMD10 (Figure 1A) on regulated recycling of ENaC, endogenous amiloride-sensitive ENaC short circuit current (*Isc*) was measured in mCCDcl1 control or COMMD10 knockdown epithelia using forskolin (FSK) stimulation. Figure 1B shows a representative trace of ENaC *Isc* and trans-epithelial resistance where due to the intensified ENaC insertion to the PM by FSK stimulation, the increase in *I*sc was accompanied by simultaneous reduction in trans-epithelial resistance (Figure 1B).

**Figure 1.**
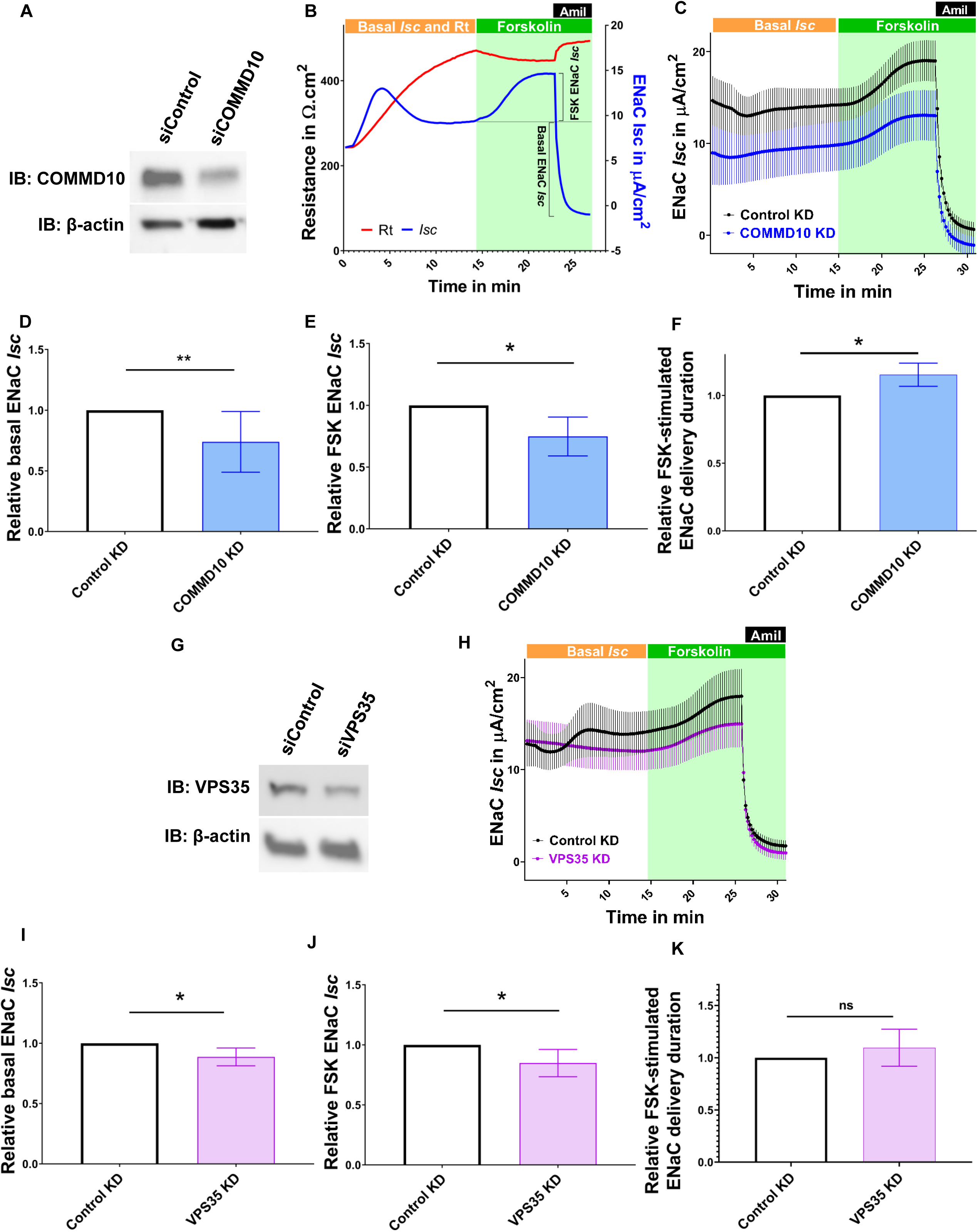
COMMD10 KD reduces regulated recycling of endogenous ENaC. (A) Representative western blots of four individual experiments reflecting COMMD10 protein levels in siControl and siCOMMD10 transfected cells. mCCDcl1 cells were transfected with either control siRNA or COMMD10 siRNA and grown for electrophysiological studies. 120-hours post-transfection, the epithelia were used in Ussing assay following which they were immediately lysed and analyzed by Western blotting with anti-COMMD10 antibody. (B) A representative trace of ENaC *I*sc and transepithelial resistance that demonstrate changes with FSK stimulation and amiloride addition. mCCDcl1 cells were transiently transfected with control siRNA or COMMD10 siRNA. 120-hours post-transfection, the *Isc* was measured and upon *Isc* stabilization, 5 μM forskolin was added basolaterally. (C) Trace of relative average ENaC *Isc* in control and COMMD10 KD epithelia (N=3, n=5, mean ± SEM). (D) Pooled basal ENaC *Isc* results showing significant reduction in COMMD10 KD epithelia. *Isc* in control KD epithelia was normalized to 1 and that in COMMD10 KD epithelia was compared to the normalized control. One-sample *t*-test; **P<0.01, mean ± SD. N=7, n=11. (E) Pooled results showing that FSK-stimulated ENaC *Isc* was reduced significantly in COMMD10 KD epithelia. *Isc* in control KD epithelia was normalized to 1 and that in COMMD10 KD epithelia was compared to the normalized control. One-sample *t*-test, *P<0.05, mean ± SD. N=3, n=5. (F) Relative normalized FSK-stimulated ENaC delivery durations. One-sample *t*-test *P<0.05, mean ± SD. N=3, n=5 (G) Representative western blots reflecting VPS35 protein levels in siControl and siVPS35 transfected cells. mCCDcl1 cells were transfected with either control siRNA or VPS35 siRNA and grown for electrophysiological studies. 120-hours post-transfection, the epithelia were used in Ussing assays following which they were immediately lysed and analyzed by Western blotting using anti-VPS35 antibody. (H) Traces of relative average ENaC *Isc* in control and VPS35 KD epithelia with FSK stimulation (N=4, n=6, mean ± SEM). mCCDcl1 cells were transiently transfected with control siRNA or VPS35 siRNA. At 120 h post-transfection, the *Isc* was measured with addition of 5 μM forskolin basolaterally. (I) Pooled results of basal ENaC *Isc* that was reduced significantly in VPS35 KD epithelia. One-sample *t*-test, *P<0.05, mean ± SD. N=4, n=6. (J) Pooled results of FSK stimulated ENaC *Isc* that was reduced significantly VPS35 KD epithelia. One-sample *t*-test, *P<0.05, mean ± SD. N=4, n=6. (K) Pooled results of FSK-stimulated ENaC delivery duration. One-sample *t*-test, P>0.05, mean ± SD. N=4, n=6. See also Figure S1 and S2.

Figure 1C demonstrates relative average traces of basal and FSK-stimulated ENaC *Isc* in control and COMMD10 KD epithelia across five experiments. Analysis of basal ENaC *Isc* in mCCDcl1 control or COMMD10 KD epithelia showed up to a ∼26% reduction in *Isc* in COMMD10 KD epithelia (1 for control KD and 0.74 ± 0.25 for COMMD10 KD epithelia; Figure 1D). Similarly, the FSK-stimulated ENaC *I*sc in COMMD10 KD epithelia was also reduced (∼ 26%) compared to control KD epithelia (1 for control vs 0.74±0.14 for COMMD10 KD epithelia; Figure 1E) where COMMD10 protein levels were suppressed by 55% (Figure 1A). As Figure 1C demonstrates, the average FSK-stimulated ENaC *Isc* trace has slid down upon amiloride addition in COMMD10 KD epithelia compared to control KD epithelia suggesting that COMMD10 KD reduces other currents as well, however, this reduction is negligible (Figure 2SA and B).

The FSK-stimulated ENaC delivery to the PM was slowed in COMMD10 KD cells (Figure 1F). Interestingly, in COMMD10 KD epithelia (compared to control epithelia), the relative reduction in basal ENaC *Isc* (1: 0.74, Figure 1D) is very similar to the relative reduction in FSK-stimulated ENaC *Isc* (1: 0.74, Figure 1E). This suggests that impairment in FSK-stimulated ENaC apical delivery, is alone responsible for the decrease in basal *Isc* in COMMD10 KD epithelia. As COMMD10 is known to function in endosomal sorting (as part of the CCC), the results suggest a role for FSK stimulation in sequence-dependent sorting of ENaC like for β2ARs, however, COMMD proteins were also shown to be involved in trafficking of recycling endosomes (Campion *et al*., 2018).

To confirm that FSK-stimulation and also thereby COMMD10 functions on the regulated recycling pathway, endogenous ENaC *Isc* was measured in mCCDcl1 epithelia with knockdown on VPS35 which is a subunit of retromer (a cargo sorting complex for regulated recycling (Cullen & Steinberg, 2018; Simonetti & Cullen, 2019)). Figure 1H demonstrates relative average traces of basal and relative FSK-stimulated ENaC *Isc* in control and VPS35 KD epithelia. With ∼35% mean decrease in VPS35 protein levels (Figure 1G), the basal ENaC *Isc* was reduced (∼12%, Figure 1I) consistent with our previous studies (McDonald lab, unpublished work). FSK-stimulated ENaC *I*sc in VPS35 KD epithelia was also reduced (∼ 15%) compared to that in control KD epithelia (1 for control vs 0.85±0.11 for VPS35 KD epithelia; Figure 1H).

The reduction in basal *Isc* in VPS35 KD epithelia (compared to control KD epithelia, 1: 0.88) is similar to the reduction in FSK-stimulated *Isc* (1: 0.85) in VPS35 KD epithelia suggesting that impairment in FSK-stimulated ENaC apical delivery is alone responsible for the decrease in basal *Isc* in VPS35 KD epithelia. The FSK-stimulated ENaC delivery to the PM was slowed in VPS35 KD cells (Figure 1K).

### 2. COMMD10 localizes to tubule-like structures emanating from Rab5- and Rab7-positive endosomes

The localization pattern of COMMD10 was evaluated on Rab5-positive endosomes in COMMD10 KD U2OS and FRT cells to identify whether COMMD10 protein localizes on the base of recycling tubules like the WASH and Arp2/3 complexes, cortactin, endosomal F-actin, and coronin (Puthenveedu *et al*., 2010), or whether COMMD10 coats the whole tubule like VPS35 (Kovtun *et al*., 2018). The results showed that COMMD10-GFP proteins are mostly concentrated on tubule-like structures emanating from endosomes (Figure 2A and C, Video S1) rather than on the whole membrane of the Rab5-positive endosome. This localization profile of COMMD10 is close to that described for EGF labelled tubules on EEA1-positive endosomes (Skjeldal *et al*., 2012). However, the COMMD10-marking on tubules are shorter than those described by Skjeldal et al. (2012) suggesting that COMMD10 also localizes to the base of tubules emanating from endosomes like endosomal F-actin, coronin, and the WASH complex (Puthenveedu *et al*., 2010) that function in concert with the CCC complex.

**Figure 2.**
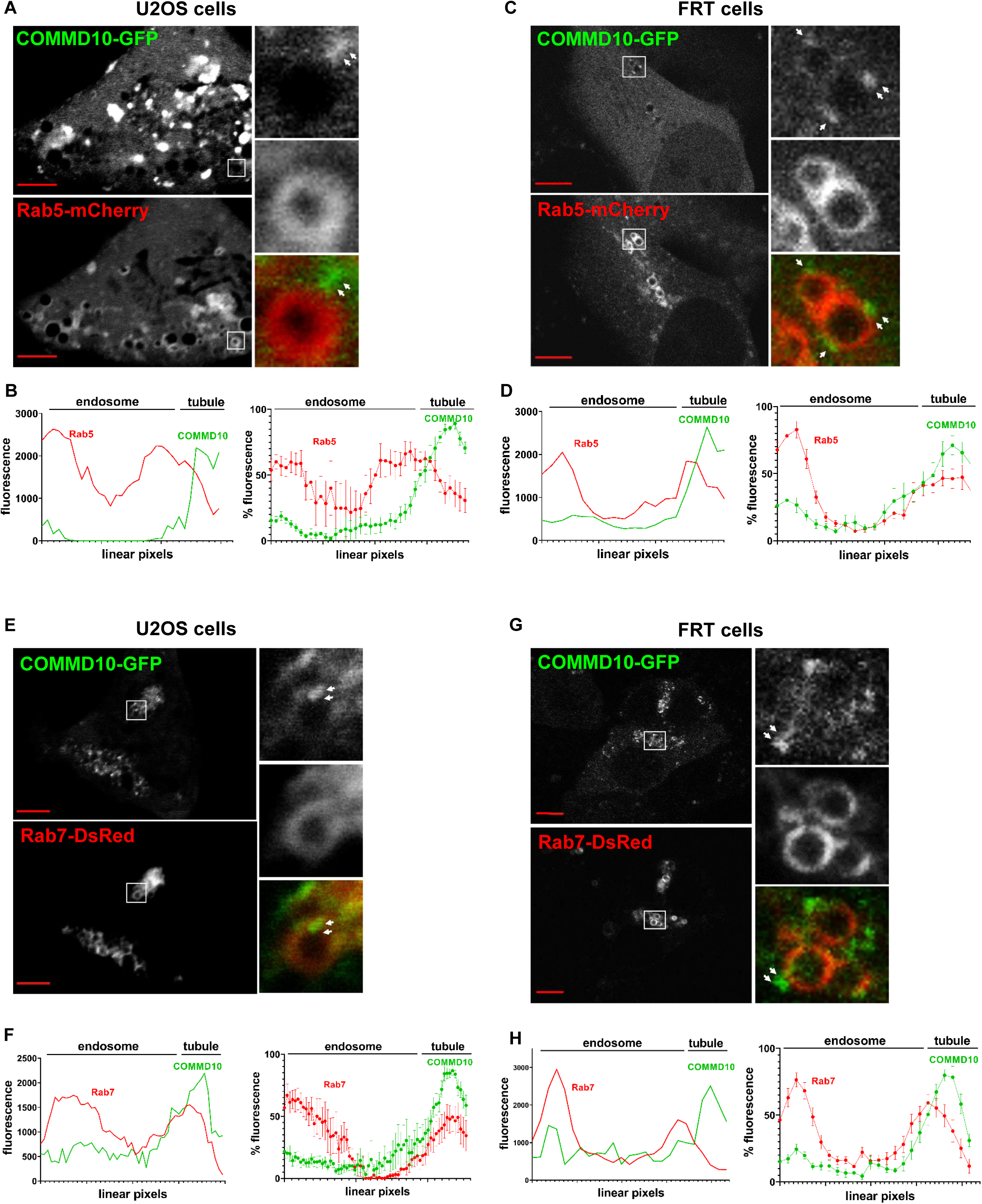
COMMD10 localizes to tubule-like structures emanating from Rab5-and Rab7-positive endosomes in U2OS and FRT cells. (A) and (C) Representative image for COMMD10 localization pattern in U2OS and FRT stable COMMD10 KD cells co-expressing COMMD10-GFP and Rab5a-mCherry, imaged by Nikon A1R, 60× PlanApo or by Olympus FV3000, 60x (1.49 NA, oil immersion), respectively. Boxed areas are enlarged in the inset where the arrows indicate concentration of COMMD10 on endosomal tubular domains. (B) and (D) - left panels. Trace of linear pixel values for Rab5 (red) and COMMD10 (green) across the same endosomes given in panels (A) and (C), confirming that the tubule/base of tubule emanating from the Rab5 endosome is enriched with COMMD10, in U2OS and FRT cells, respectively. (B) and (D) - right panels. Trace of linear pixel values for Rab5 (red) and COMMD10 (green) on endosomal tubules averaged across 12 and 16 endosomes in U2OS or FRT cells, respectively, normalized to the maximum, supporting the visualizations showing specific enrichment of COMMD10 on tubular structures. Linear regression, mean ± SEM. N=3, n=12 or n=16. (E) and (G) Representative image for COMMD10 localization pattern in U2OS and FRT stable COMMD10 KD cells co-expressing COMMD10-GFP and Rab7a-DsRed, imaged by Nikon A1R, 60× PlanApo or by Olympus FV3000 (1.49 NA, oil immersion), respectively. The boxed areas are enlarged in the images on the right panel where the arrows indicate concentration of COMMD10 on a Rab7-positive endosomal tubular domain. (F) and (H) - left panels. Trace of linear pixel values for Rab7 (red) and COMMD10 (green) across the same endosomes given in panels (E) and (G), confirming that the specific domain on the Rab7-positive endosome is enriched with COMMD10. (F) and (H) - right panels. Trace of linear pixel values for Rab7 (red) and COMMD10 (green) on endosomal tubules averaged across 11 and 16 endosomes in U2OS or FRT cells, respectively, normalized to the maximum, which supports the specific enrichment of COMMD10 on endosomal domains. Linear regression, mean ± SEM. N=4, n=11 or n=16. See also Video S1.

Linear pixel analysis of fluorescence intensity along the endosome also shows that COMMD10 is not recruited onto the whole Rab5-positive endosome but only to the tubular structure emanating from the endosome (Figure 2B and D). The fluorescence intensity trace in the right panel of Figure 2B and D were obtained from the same endosomes shown in Figure 2A and C while the average traces in the right panel of Figure 2B and D are from 12 and 16 individual endosomes, respectively.

Retromer-mediated endosomal sorting continues on Rab7-positive endosomes (Purushothaman *et al*., 2017) which still requires the WASH and CCC complex activity for tubule release. To examine the localization of COMMD10 on microdomains of Rab7-positive endosomes, COMMD10-GFP and Rab7a-DsRed were coexpressed in U2OS and FRT COMMD10 stable knockdown cells and evaluated by confocal microscopy.

The results show similar COMMD10 localization pattern on tubular structures of Rab7-positive endosomes like that on the Rab5-positive endosomes (Figure 2E and G). The linear pixel analysis of fluorescence intensity along the endosomes confirm that COMMD10 recruitment onto specific tubule-like structures on Rab7-positive endosomes continues (Figure 2F and H, left panels), possibly until the end of cargo sorting. The average traces in (Figure 2F and H, right panels) are from 11 and 16 individual endosomes, respectively in U2OS and FRT cells.

The results from both the U2OS and FRT cell lines show that COMMD10 localizes onto specific domains associated with Rab5-and Rab7-positive endosomes supporting the possibility of these domains are sequence-dependent endosomal sorting microdomains consistent with the hypothesis that COMMD10 as a part of CCC complex having a role in sorting of ENaC at this location.

### 3. COMMD10 migrates on Rab11-positive recycling endosomes

COMMD5 was shown to hook recycling endosomes to the cytoskeleton (Campion *et al*., 2018) suggesting a role for COMMDs in translocation of recycling vesicles. To evaluate this hypothesis with COMMD10, COMMD10-GFP and Rab11a-DsRed were coexpressed in U2OS and FRT stable COMMD10 knockdown cells and COMMD10 colocalization with Rab-11 positive recycling endosomes was evaluated. In the U2OS cells, Rab11-positive recycling endosomes were observed in the peripheral regions of the cells, particularly around the cellular tips (edges) (Figure 3A), consistent with previous reports (Bruce *et al*., 2012; Takahashi *et al*., 2012), but inconsistent with many other reports showing the Rab11-positive structures concentrated to the perinuclear region as an ERC (Horgan *et al*., 2010; Amorim *et al*., 2011; Mölleken & Hegemann, 2017).

**Figure 3.**
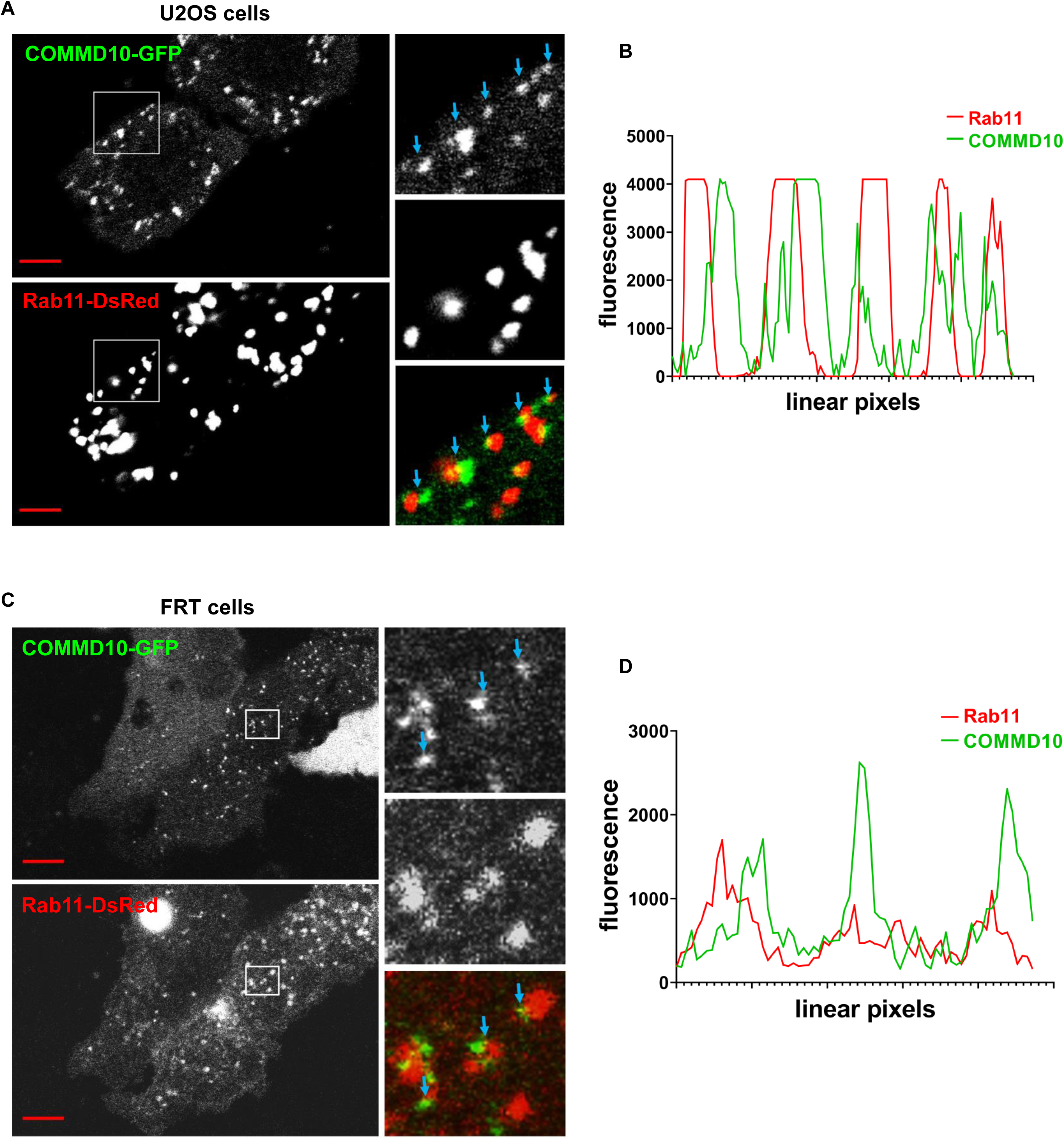
COMMD10 concentrates on specific structures on Rab11-positive recycling endosomes. (A) Representative image for COMMD10 localization pattern in U2OS stable COMMD10 KD cells coexpressing Rab11a-DsRed and COMMD10-GFP, imaged by Nikon A1R, 60× PlanApo (1.49 NA, oil immersion). Boxed areas are enlarged in the inset where the arrows indicate concentration of COMMD10 on endosomal tubular domains. (B) Trace of linear pixel values across the same row of endosomes, confirms that COMMD10 is enriched in specific structures on Rab11-positive recycling endosomes. N=3. (C) Representative image for COMMD10 localization pattern in FRT stable COMMD10 KD cells coexpressing Rab11a-DsRed and COMMD10-GFP, imaged by Olympus FV3000, 60x (1.49 NA). Boxed areas are enlarged in the inset where the arrows indicate concentration of COMMD10 on endosomal tubular domains. (D) Trace of linear pixel values across the same row of endosomes, confirms that COMMD10 is enriched in specific structures on Rab11-positive recycling endosomes. N=2. See also Video S2.

COMMD10 was concentrated on most of the Rab11-positive endosomes (Figure 3A) but in an asymmetric pattern, with occasional areas of overlap (blue arrows). In the time-lapse series of images, COMMD10- and Rab11-positive vesicular structures are seen to move in an intermittent manner (Video S2) suggestive of microtubule-based transport consistent with previous reports (Amorim *et al*., 2011; Takahashi *et al*., 2012). Linear pixel analysis of fluorescence intensity along the Rab11-positive recycling endosomes (Figure 3B), obtained from the same row of recycling endosomes shown in the Figure 3A, confirms that COMMD10 is not recruited onto the whole membrane of Rab11-positive endosome but localizes to specific structures on the endosomes. Consistent results are seen with the COMMD10 localization pattern on Rab11-positive endosomes of FRT cells (Figure 3C and D) suggesting that COMMD10 may have a role in translocation of recycling endosomes to the PM.

### 4. Aldosterone reduces COMMD protein levels, but not mRNA levels, in early phase of action

In search of regulators of COMMD proteins and ENaC recycling pathways, we evaluated the effect of aldosterone on COMMDs as aldosterone is known as the main upregulator of ENaC in the kidney. After incubation of mCCDcl1 cells with aldosterone for various time periods, ENaC *Isc* was measured in this epithelia where the results confirmed that aldosterone significantly increases ENaC Isc in mCCDcl1 epithelia. Thus, aldosterone increased ENaC *Isc* at all time periods (0.5 hour: 1 for control vs 1.57 ± 0.22 for aldo-treated epithelia; 3 hours: 1 for control vs 1.34 ± 0.34 for aldo-treated epithelia; 24 hours: 1 for control vs 2.419 ± 0.652 for aldo-treated epithelia, Figure 4A). Then, COMMD1 and −10 protein levels were analyzed with the same way, treating the mCCDcl1 cells with aldosterone for the time periods (Figure 4B). COMMD1 and −10 protein levels were significantly decreased after 300 nM aldosterone treatment for 30 min (COMMD1: 1 for vehicle vs 0.63±0.09 fo0r aldo; COMMD10: 1 for vehicle vs 0.76 ±0.09 for aldo) but not after longer treatment periods (Figure 4D and E).

**Figure 4.**
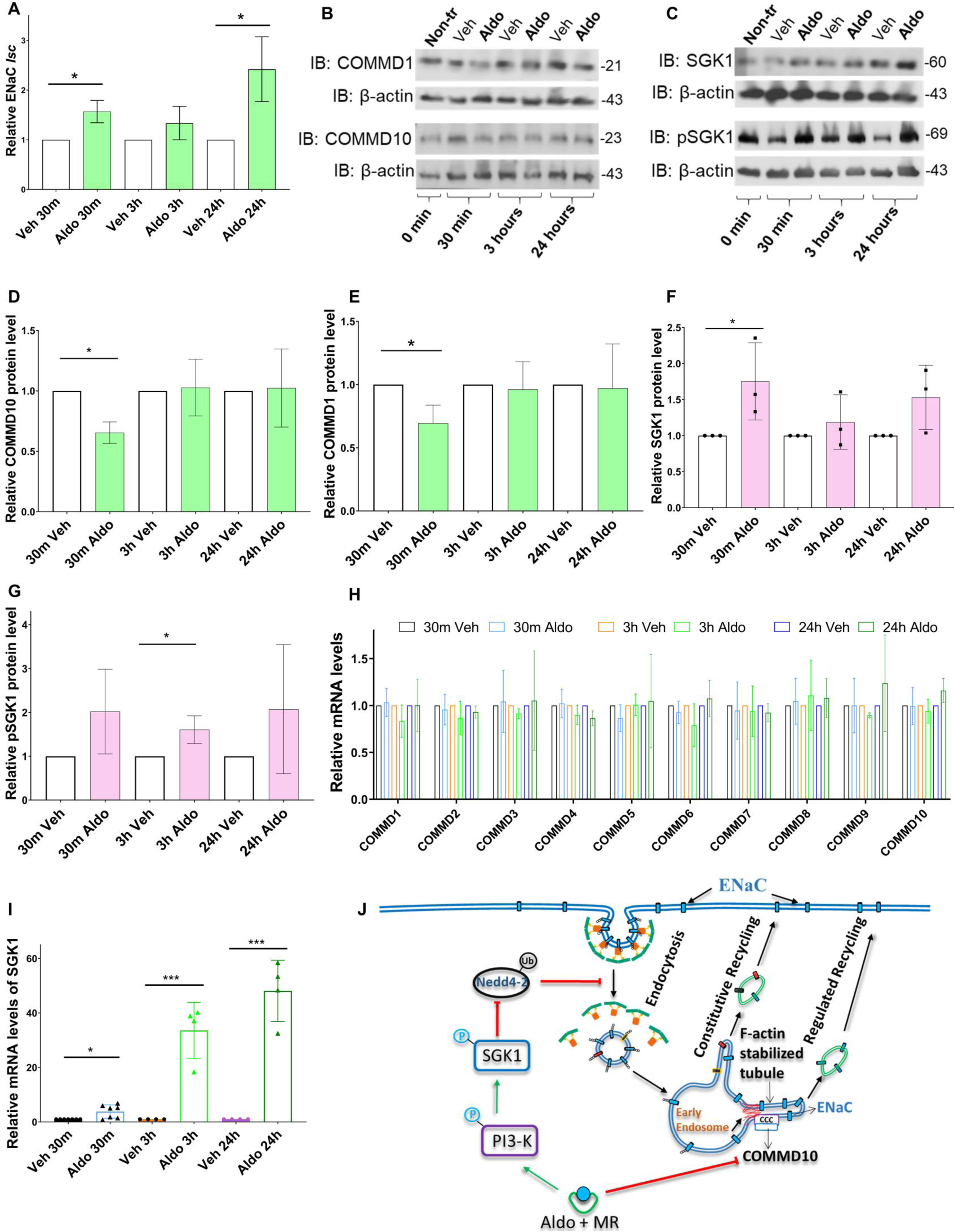
Aldosterone reduces COMMD protein levels, but not mRNA levels, in early phase of action. (A) Pooled relative ENaC *Isc* results showing significant increase with 0.5- and 24-hour aldosterone treatment. mCCDcl1cells, grown on permeable membranes, were treated with aldosterone (Aldo, 300 nM) or vehicle (Veh, 100% DMSO) 144 hours after seeding the cells for 0.5, 3, and 24 hours. 0.5 hour: *P<0.05; 3hours: P=0.09; 24hours: *P<0.05. One-sample *t*-test for each veh-aldo pair separately, mean ± SD. N = 4. (B) Representative blots for COMMD1 and −10 protein levels in mCCDcl1 cells, respectively. mCCDcl1 cells were grown on permeable membranes, treated with 300 nM aldosterone or vehicle for 0.5, 3, and 24 h in serum and drug free medium. (C and D) Pooled relative protein levels of COMMD1 and −10 that were decreased with only 30 min aldo-treatment. One-sample t-test for each veh-aldo pair. *P<0.05, mean ± SD. N=4 for COMMD1, N=3 for COMMD10. (E) mRNA levels of COMMD1-10 showing no change of significance (RQ values were not beyond 0.5 and 2) after 300 nM aldosterone treatment for 0, 0.5, 3, and 24-hour time periods. One-way ANOVA for each COMMD separately. N=7 for 30 min, N=4 for 3 and 24 hour periods. (I) mRNA levels of SGK1, as a positive control for aldosterone treatment, were increased significantly at all three time periods. *Veh*-vehicle (DMSO), *Aldo*-aldosterone (in DMSO). One-sample *t*-test data. N=7 for 30 min, N=4 for 3 and 24 hour periods. *P˂0.05, **P˂0.01. (J) Schematic model for early phase of aldosterone effect on ENaC trafficking and COMMD proteins. See also Figure S3

COMMD1-10 genes are expressed in various levels (Figure S3A) as protein abundance have been shown to vary in mouse mpkCCD cells (Yang *et al*., 2015) and the variance in mRNA and protein levels do not overlap (Figure S3). Analysis of mRNA levels of COMMDs in 300 nM aldosterone treated cells revealed that aldosterone does not change the mRNA levels of any of COMMD1-10 (RQ values were not beyond 0.5 and 2) at any time period (Figure 4I) suggesting the reduction in protein levels of COMMDs by aldosterone is not linked with changes in mRNA levels. However, SGK1 mRNA levels (a positive control for aldosterone treatment) were significantly increased at each time period (30 min: 1 for vehicle vs 3.82±2.45 for aldo-treated epithelia; 3 hours: 1 for vehicle vs 33.58±10.28 for aldo-treated epithelia; 24 hours: 1 for vehicle vs 48.12±11.23 for aldo-treated epithelia, Figure 4J) consistent with previous reports (Náray-Fejes-Tóth *et al*., 1999; Jacobs *et al*., 2016; Welch *et al*., 2016).

### 5. Calcium treatment upregulates COMMD protein levels and induces basal ENaC *Isc*

COMMD5 was shown to be negatively regulated by extracellular calcium treatment, and its basal mRNA and protein levels were found to be higher in hypertensive animals (Solban *et al*., 2000). Therefore, it was hypothesized that COMMDs in principal cells of distal nephron can be regulated by extracellular calcium and thereby can modulate ENaC cell surface activity. To test this hypothesis, mCCDcl1 cells were treated with extra Ca^2+^ and protein levels of COMMD1 and COMMD10 and also mRNA levels of COMMD1, COMMD10, and COMMD5, were analyzed. Surprisingly, the results showed that COMMD10 protein levels were increased (also an increasing trend in COMMD1 level followed) with an extra 1, 2, or 4 mM CaCl_2_ treatment (but not with 6 mM) for a total of 72 hours (Figure 5A, B, C). Further analysis showed that, however, the increase in protein levels of COMMDs by Ca^2+^ treatment is not linked with changes in mRNA levels of COMMDs after treating mCCDcl1 cells with an extra 2 and 4 mM of CaCl_2_ (Figure 5D).

**Figure 5.**
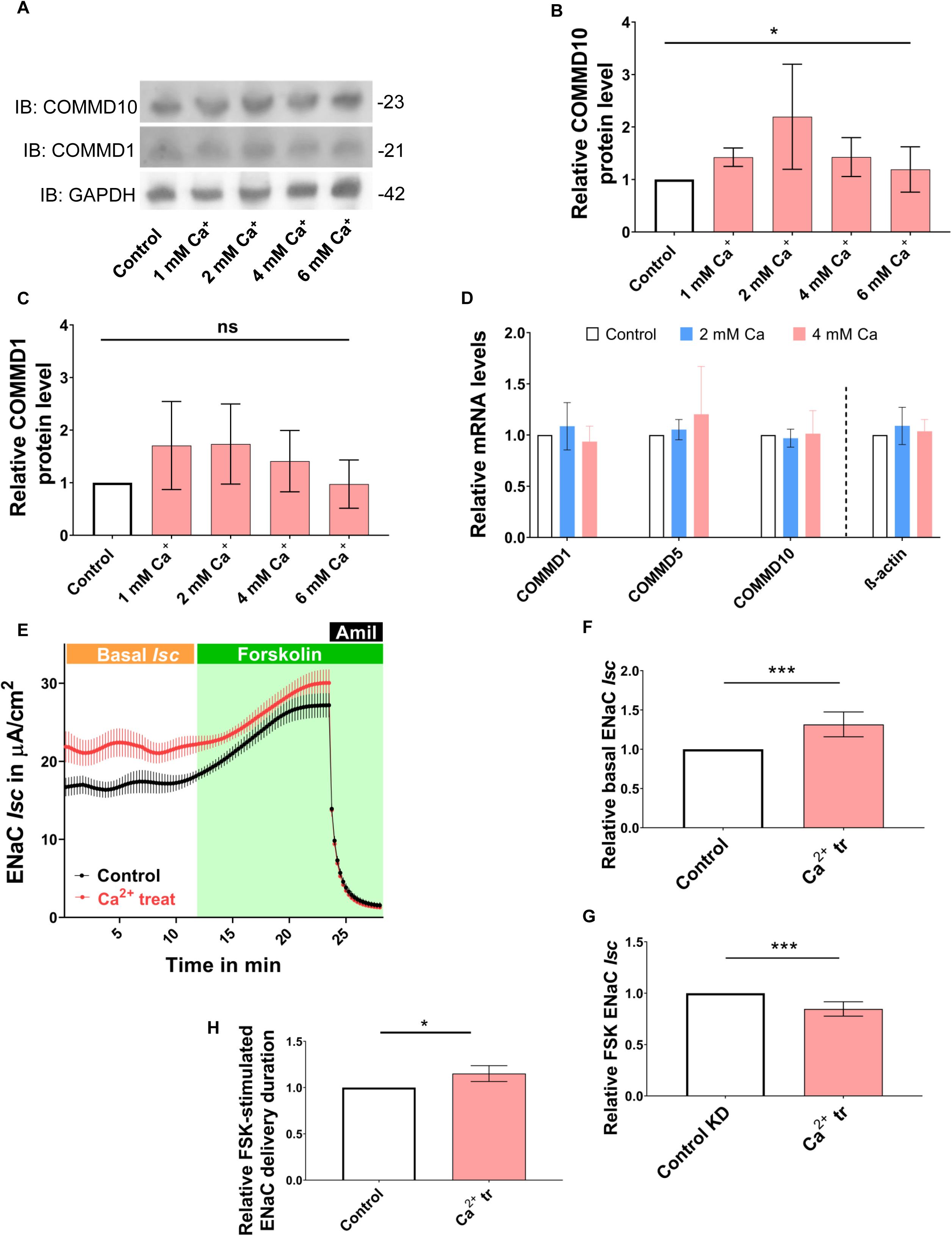
Ca^2+^ treatment stimulates COMMD1, and −10 protein levels but doesn’t alter mRNA levels of COMMD1, 5, and 10 in mCCDcl1 cells. (A) Representative blots of COMMD1 and −10 including GAPDH as loading control. mCCDcl1 cells were treated with an extra 1, 2, 4, or 6 mM CaCl_2_ for a total of 72 hours and analyzed by Western blotting. (B and C) Pooled relative protein levels of COMMD1, and −10. One-way ANOVA, *P˂0.05, means ± SD. N=4 or N=3. (D) mRNA levels of COMMD1, −5 and −10 that were not altered by Ca^2+^ treatment. mCCDcl1 cells were grown on permeable membranes, treated with 2 or 4 mM CaCl_2_ for a total of 72 hours. mRNA levels of reference gene β-actin were also not altered by extra Ca^2+^ that analyzed by normalizing Ct values of β-actin in control cells to one and comparing the Ct values of β-actin in treated cells to the normalized control. P>0.05, N=3. (E) Trace of relative average ENaC *Isc* in control and Ca^2+^ treated epithelia demonstrating relative changes with FSK stimulation (N=4, n=8, mean ± SEM). mCCDcl1 cells were treated with an extra 2 mM CaCl_2_ for a total of 96-hours. The *Isc* was measured and upon *Isc*-stabilization, the measurement was continued with addition of 5 μM forskolin basolaterally. (F) Basal ENaC *Isc* was increased by Ca^2+^ treatment. One-sample t-test, ***P<0.001, mean ± SD. N=4, n=8. (G) FSK stimulated ENaC *Isc* was reduced. One-sample t-test, ***P<0.001, mean ± SD. N=4, n=8. (H) Pooled results of FSK-stimulated ENaC delivery durations. One-sample *t*-test, *P<0.05, mean ± SD. N=4, n=8.

To test how regulated ENaC cell surface population by Ca^2+^, endogenous ENaC *Isc* was measured after treating the epithelia with 2 mM excess Ca^2+^ (a total of 3.5 – 4 mM Ca^2+^) for a total of 96 hours. Also to understand the effect of Ca^2+^ treatment on ENaC recycling, the *Isc* measurement was continued with 5 μM (final concentration) FSK treatment upon the *Isc* stabilization. Figure 5E demonstrates relative average traces of basal and FSK-stimulated ENaC *Isc* in control and Ca^2+^ treated epithelia across four individual experiments. Basal ENaC *Isc* in Ca^2+^ treated mCCDcl1 epithelia was increased up to a 31% compared to control epithelia (1 for control and 1.31 ± 0.17 for Ca^2+^ treated epithelia, Figure 5F). However, interestingly, FSK-stimulated ENaC *Isc* in Ca^2+^ treated epithelia was (15%) less than that in control epithelia (1 for control vs 0.85±0.07 for Ca^2+^ treated epithelia, Figure 5G).

Furthermore, the FSK-stimulated ENaC apical delivery duration was also reduced (which means recycling fastened) in Ca^2+^ treated epithelia compared to control epithelia (1 for control, 0.87±0.12 for Ca^2+^ treated epithelia, Figure 5H). This suggests that the increase in COMMD10 levels by Ca^2+^ treatment is not a result of compensation for their functional blockage on endosomal sorting (unlike COMMD10 KD which slowed recycling, thereby increased ENaC apical delivery duration).

### 6. Calcium treatment leads to microfilament reorganization and endosomal F-actin accumulation

A study demonstrated that 5 mM Ca^2+^ treatment inhibited FSK-stimulated AQP2 levels at the apical membrane of collecting duct cells, identifying twofold increase in F-actin content as one of the pathways contributing to this effect (Procino *et al*., 2004). COMMD5 KD has also been shown to alter microfilament organization (Campion *et al*., 2018). Linking these two results, we evaluated microfilament organization in Ca^2+^-treated cells and also in COMMD10 KD cells. The results showed that COMMD10 KD doesn’t alter microfilament organization in U2OS (Figure 6A and B) and FRT cells (Figure S3), however, qualitative analysis show that Ca^2+^-treatment reorganizes microfilament distribution and reduces fluorescence intensity of microfilaments in U2OS cells where it is expected that COMMD10 levels are higher. Thus, the F-actin stress fibers demonstrate a more disordered distribution and its fluorescence intensity was significantly reduced in Ca^2+^ treated cells compared to the control cells (1170±480 for control cells vs 780±250 for Ca^2+^ treated cells, Figure 6A lowest panel and Figure 6C).

**Figure 6.**
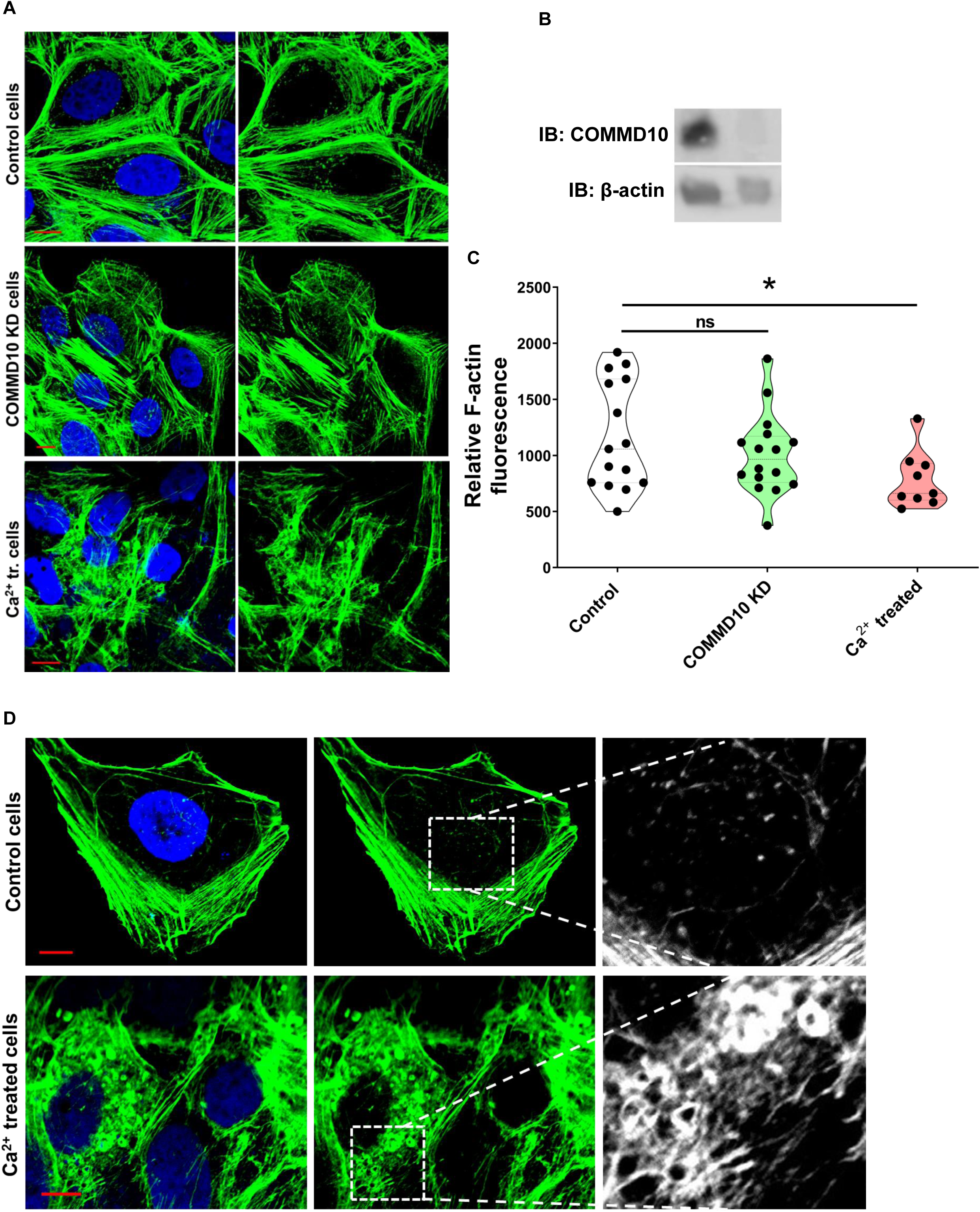
Calcium treatment leads to microfilament reorganization and endosomal F-actin accumulation. (A) Representative images of stained F-actin in U2OS control, COMMD10 KD, and Ca^2+^ treated cells (with an extra 2 mM CaCl_2_. (B) Representative western blots for COMMD10 KD in U2OS cells. (C) Quantitative analysis of fluorescence intensity of actin filaments in control, COMMD10 KD, and Ca^2+^ treated cells showing less F-actin intensity in Ca^2+^ treated cells compared to control cells. Student’s t-test, *P<0.05. N=2, n=8, representing 117 (control), 153 (COMMD10 KD), and 92 (Ca^2+^ treated) cells. (D) Ca^2+^ treatment (2 mM extra) also caused endosomal F-actin accumulation and vesicle enlargement in U2OS cells. Scale bar = 10 μM.

As seen in the figures of both U2OS and FRT cells (Figure 6A and Figure S3), some special types of vesicles were stained by phalloidin together with F-actin stress fibers. Surprisingly, more detailed analysis showed that in the Ca^2+^ treated U2OS cells, F-actin was accumulated on enlarged vesicles compared to those in control cells (Figure 6D) suggesting endosomal actin nucleation activity was increased, as no other types of vesicles are known to be covered by actin except endosomes and also recycling/secretory vesicles but after fusion with the PM (Miklavc *et al*., 2012).

## Discussion

The activity of ENaC in the apical membrane determines the extent of Na^+^ reabsorption across an epithelium. For downregulation of Na^+^ reabsorption, ENaC is removed from the apical membrane through internalization (Shimkets *et al*., 1997; Wang *et al*., 2006; Wiemuth *et al*., 2007), while for upregulation, ENaC internalization is prevented (Debonneville *et al*., 2001; Snyder *et al*., 2002; Balderhaar *et al*., 2010; Hallows *et al*., 2010; Lang & Stournaras, 2013), and biosynthetic pathway (Girardet *et al*., 1986; Loffing *et al*., 2001) (Asher *et al*., 1996; Gaeggeler *et al*., 2005) or recycling pathways are activated (Butterworth *et al*., 2009; Butterworth *et al*., 2012; Edinger *et al*., 2012; Klemens *et al*., 2017). The regulated recycling pathway involves multiple protein complexes of endosomal sorting machinery including SNXs, WASH/Arp2/3 complex, CCC complex, retromer and retriever complexes etc. Internalized ENaC is delivered to early endosomes where ENaC destined for recycling back to the plasma membrane is sorted into a pathway distinct from proteins that are sent for lysosomal degradation. However, it is not known if ENaC is sorted for recycling by the conventional sorting machines of WASH/CCC axis.

Burstein and colleagues showed that COMMD family proteins are part of the CCC complex (Starokadomskyy *et al*., 2013) that is recruited onto endosomes through interaction with FAM21 and the retromer complex (Phillips-Krawczak *et al*., 2015). The CCC complex is also very important for the recruitment of another multiprotein complex, retriever, that functions in endosomal sorting like retromer (McNally *et al*., 2017). The endosomal-recruited COMMDs, as a part of the CCC complex, regulate recycling of many membrane proteins such as ATP7A, Na-K-2Cl cotransporter, CFTR, LDL-R, Notch signaling receptor etc. (Drevillon *et al*., 2011; Smith *et al*., 2013; Li *et al*., 2015; Phillips-Krawczak *et al*., 2015; Bartuzi *et al*., 2016; McNally *et al*., 2017). Although COMMD5 was shown to regulate EGFR by hooking recycling endosomes to the cytoskeleton, exact role of other COMMDs, in endosomal sorting is unknown. Also up to now, no specific regulators for COMMD proteins were identified. In the present study, we found that: (1) COMMD10 KD slows endocytic recycling of ENaC, (2) COMMD10 localizes to distinct structures on Rab5- and Rab7-positive endosomes and on Rab11-positive recycling vesicles, (3) COMMD1 and −10 protein levels are reduced by aldosterone in its early phase of action, (4) extracellular calcium treatment leads to endosomal F-actin accumulation, increase in COMMD1 and −10 protein levels, and enhanced ENaC *Isc*.

### COMMD10 knockdown impairs regulated recycling of ENaC

ENaC is recycled to the cell surface via constitutive recycling and separately via a cAMP-regulated recycling pathways (Butterworth *et al*., 2009; Butterworth *et al*., 2012; Edinger *et al*., 2012; Klemens *et al*., 2017) like β2AR (Puthenveedu *et al*., 2010; Bowman *et al*., 2016). Upon cAMP stimulation, ENaC is recruited from intracellular stores and inserted into the apical membrane (Kleyman *et al*., 1994; Snyder, 2000; Morris & Schafer, 2002), while further investigation showed that this insertion is through translocation of subapical vesicles (Butterworth *et al*., 2005; Edinger *et al*., 2012) suggesting cAMP stimulation upregulates ENaC sorting to the regulated recycling tubules/vesicles as it does the β2AR (Vistein & Puthenveedu, 2013; Bowman *et al*., 2016).

In many studies, suppression of endosomal sorting machinery subunits revealed accumulation of cargoes on endosomes (Liu *et al*., 2012a; Liu & Grant, 2015; Norris *et al*., 2017; Singla *et al*., 2019). However, the mechanism of cargo trapping in endosomes is not known. We have previously reported that ENaC *Isc* is reduced in COMMD10 KD cells due to the loss of ENaC population at the cell surface (Ware *et al*., 2018) and here we demonstrate that the decrease in the ENaC population at the cell surface of COMMD10 KD cells is not the result of fast ENaC endocytosis but an accumulation of ENaC intracellularly supporting a role for COMMD10 in ENaC endosomal sorting or recycling. Furthermore, using forskolin as a cAMP stimulator (Garty & Palmer, 1997; Morris & Schafer, 2002; Snyder *et al*., 2004; Butterworth *et al*., 2005), we showed that COMMD10 KD reduces FSK-stimulated endogenous ENaC *Isc* as well as basal ENaC *Isc* in mCCDcl1 epithelia which confirms that COMMD10 KD impairs ENaC recycling. The role of COMMD10 could be in endosomal sorting (through regulation of WASH complex recruitment (Singla *et al*., 2019)) or in translocation of recycling vesicles (through hooking endosomes to cytoskeleton (Campion *et al*., 2018)). However, VPS35 KD also reduced FSK-stimulated ENaC *Isc* supporting the idea that COMMD10 also functions in endosomal sorting like VPS35. Secondly, the relative reduction in basal *Isc* in COMMD10 KD and VPS35 KD epithelia (compared to their control epithelia) was very similar to the relative reduction in FSK-stimulated ENaC *Isc* in these epithelia (compared to their control epithelia) suggesting that impairment in FSK-stimulated ENaC apical delivery, is alone responsible for the decrease in basal *Isc*. Therefore, it is less possible that COMMD10, like VPS35 is involved in other pathways such as constitutive recycling and biosynthetic pathways.

Furthermore, FSK-stimulated ENaC delivery duration was increased (which means recycling was slowed) in COMMD10 KD and (although no firm conclusions can be drawn because of the modest sample size) VPS35 KD epithelia compared to control epithelia. This suggests that COMMD10 KD leads to trapping of ENaC in endosomes and as long as the effect of FSK is there (whose effect lasts far longer than our experiment duration (Butterworth *et al*., 2005)), ENaC is rescued from the “trap”, but over a longer time period than in control cells, and with some loss to degradative pathway. However, these results are not sufficient to conclude that COMMD10 does not function in translocation of ENaC to the PM.

### COMMMD10 is recruited onto specific endosomal domains

The localization of WASH, Arp2/3, cortactin and coronin on endosomal domains was investigated previously (Puthenveedu *et al*., 2010), however, there have been no reports on the localization of the CCC complex or COMMD proteins on endosomal domains. Although COMMD5 colocalization with endosomal markers Rab5, −7 and −11 has been reported (Campion *et al*., 2018), the exact localization of COMMDs on endosomal domains was not evaluated. The results reported here show that COMMD10 localizes on Rab5- and Rab7-positive endosomes on specific tubule-like structures and this type of localization is similar to localization of WASH, coronin, and cortactin (Puthenveedu *et al*., 2010), and supports a recent model where the CCC complex negatively regulates WASH complex localization on endosomes (Singla *et al*., 2019).

It has been shown that the conversion of Rab5 positive endosomes to Rab7 positive ones is a gradual process (Rink *et al*., 2005) which explains why COMMD10 can localize on both early endosomes (Rab5 positive) and late endosomes (Rab7 positive). Rab7 is required for retromer recruitment onto endosomes (Rojas *et al*., 2008; Seaman *et al*., 2009; Balderhaar *et al*., 2010; Liu *et al*., 2012b; Priya *et al*., 2015) and the retromer, upon recruitment onto tubules, releases Rab7 from this domain, which separates endosomal recycling from Rab7-dependent lysosomal fusion (Purushothaman *et al*., 2017).

COMMD10 localization on Rab11-positive recycling endosomes was distinct from the localization patterns on Rab5- and Rab7-positive endosomes suggesting another role for COMMD10 in the recycling pathway, possibly similar role shown for COMMD5 to hook endosomes to cytoskeleton (Campion *et al*., 2018), but this hypothesis needs further investigation. Over-expression and knockdown of Rab11 resulted in reduced ENaC cell surface activity (Butterworth, 2010; Butterworth *et al*., 2012) suggesting Rab11-positive endosomes are recruited for ENaC recycling. Furthermore, both Rab4 and Rab11, which functions in endosome-to-plasma membrane recycling (Zerial & McBride, 2001; Maxfield & McGraw, 2004), were localized to endosomal domains containing β2AR (Puthenveedu *et al*., 2010) suggesting that after endosomal sorting, cargoes utilize both Rab4-(fast) and Rab11-mediated (slow) recycling pathways and together with cargo, COMMDs also migrate on Rab11-positive endosomes.

### COMMD proteins are regulated by aldosterone and extracellular calcium

Aldosterone, an upregulator of ENaC, that increased endogenous ENaC *Isc* in mCCDcl1 epithelia at all 0.5, 3, and 24-hour time periods of treatment, interestingly reduced COMMD1 and −10 protein levels by ∼35% at only 0.5-hour time period but not at others. Aldosterone, in its short-term effect on ENaC, prevents ENaC endocytosis through the SGK1/Nedd4-2 pathway (Balderhaar *et al*., 2010; Hallows *et al*., 2010; Lang & Stournaras, 2013). It is possible that, to preserve a constant plasma membrane (Houy *et al*., 2013; Liang *et al*., 2017), recycling is reduced by reducing the COMMD protein levels. However, possibly as ENaC-carrying membrane vesicles from the biosynthetic pathway fuse with the PM during the long-term response to aldosterone, the endocytosis pathway “opens” and the reducing effect of aldosterone on COMMD protein levels at 3- and further hours are eliminated.

As the *Hypertension-related, Calcium-Regulated Gene* (HCaRG)/COMMD5 was shown to be negatively regulated by extracellular calcium treatment (Solban *et al*., 2000), in the search for regulators of COMMD10, extracellular Ca^2+^ was found to upregulate COMMD1 and −10 protein levels in a concentration dependent manner. Ca^2+^ treatment also reduced intensity of stress fibers similar to the results for COMMD5 KD (Campion *et al*., 2018). Further analysis revealed that F-actin was accumulated on large vesicles in Ca^2+^ treated cells compared to control cells. No reports were found showing F-actin covered vesicles except endosomes. According to previous reports, endosomal F-actin accumulation occurs when the CCC complex was silenced and thereby WASH activity was increased (Singla *et al*., 2019) suggesting Ca^2+^ treatment may induce endosomal F-actin accumulation. In our case, Ca^2+^ treatment resulted in an increase in COMMD10 protein level and in basal ENaC *Isc* suggesting the possibility that CCC activity was not blocked by Ca^2+^ but the F-actin accumulation under Ca^2+^ treatment is due to intensified actin nucleation. Although there is no enough evidence to conclude this, however, it is possible that the increase in COMMD10 protein levels is due to increased CCC recruitment by WASH complex (Phillips-Krawczak *et al*., 2015) that is needed to overcome the endosomal F-actin accumulation and thereby to release recycling tubules (which is supported by the increase in ENaC *Isc* in Ca^2+^ treated cells). Surprisingly, FSK-induced ENaC *Isc* was reduced in Ca^2+^ treated epithelia compared to non-treated epithelia possibly due to depleted ENaC levels in endosomes through increased actin nucleation and COMMD10 activity. It was shown that apically increased Ca^2+^ concentration inhibits ENaC activity through intensified endocytosis of ENaC by protein kinase C (PKC) pathway (Vallon & Rieg, 2011; Toney *et al*., 2012) while basolaterally increased Ca^2+^ concentration in the cytosol very near the membrane of the cell was shown to stimulate ENaC at the cell surface of renal epithelium (Zhang *et al*., 2007; Thai *et al*., 2014).

Overexpression of COMMD10 caused cell death in many attempts preventing the direct measurement of the impact of overexpressed COMMD10 on ENaC activity in an Ussing assay. Colocalization analysis of fluorophore-labelled ENaC and COMMD10 on Rab-positive endosomes would give clearer understanding of the role of COMMD10 in ENaC recycling. A schematic model of the role of COMMD10 in ENaC endosomal sorting and recycling is described in Figure 7.

**Figure 7.**
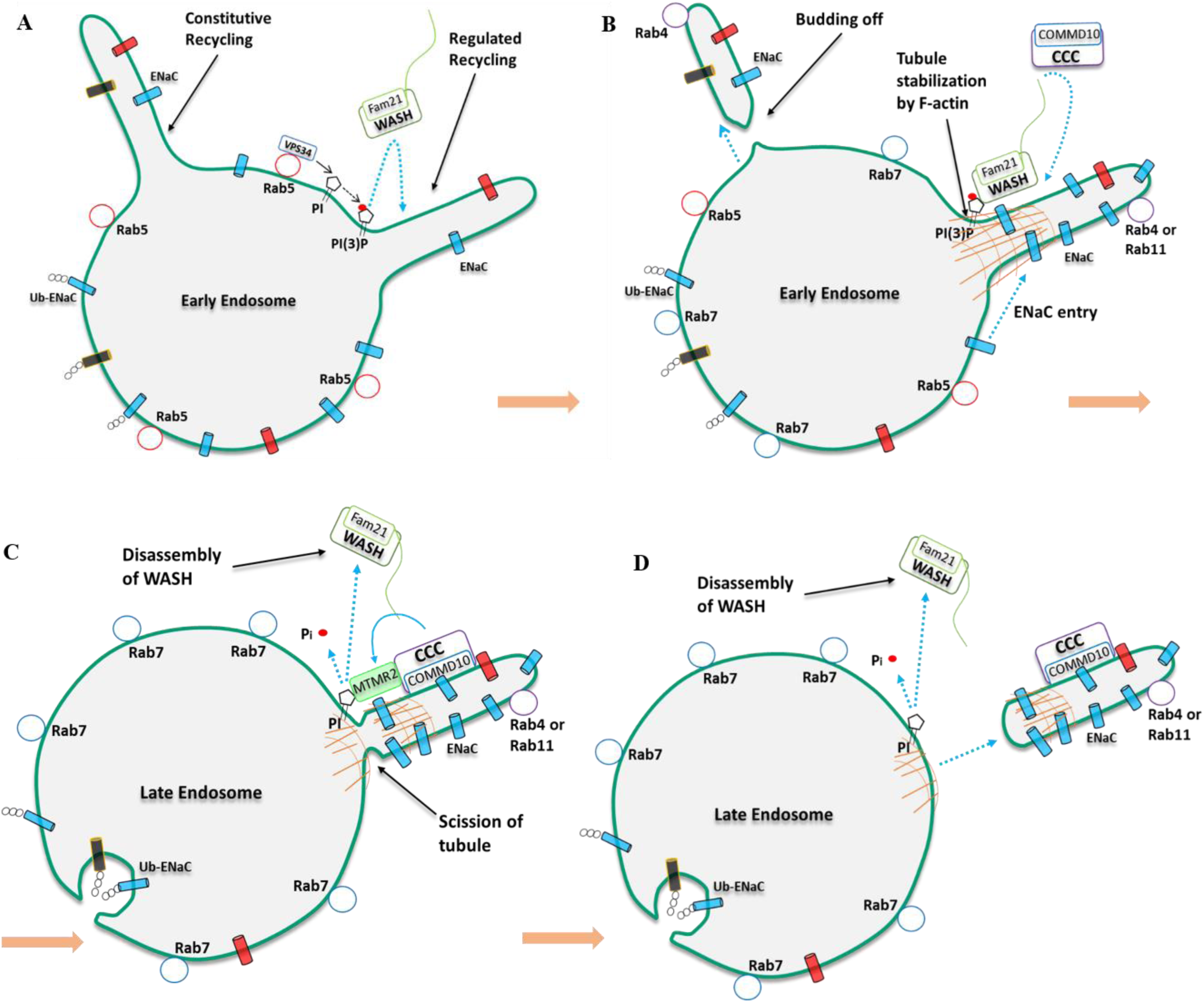
Schematic model of the role of COMMD10 in ENaC endosomal sorting and recycling. (A) Continuous generation of endosomal recycling tubules. On the base of some of those tubules, endosomal Rab5 recruits PI(3)-K named VPS34 which converts PI to PI(3)P. PI(3)P provides a base for WASH complex recruitment onto endosomes. (B) WASH complex promotes endosomal F-actin polymerization which prevents tubules from quick budding off, stabilizing them for a longer time to enable the entry of sorted cargoes. Some other tubules are released from endosomes immediately without actin-stabilization forming the bulk membrane flow (constitutive recycling) and carrying random membrane proteins and lipids such as TfR, C6-NBD-SM, and ENaC without sorting. (C) CCC complex (including COMMD10) is recruited onto the base of endosomes by the WASH complex where the CCC complex negatively regulates WASH assembly through recruiting myotubularin-related protein-2 (MTMR2) to dephosphorylate PI(3)P to PI. It is possible that as cargoes are loaded into actin-stabilized tubules, COMMD-cargo (including ENaC) interaction defines the abundance of CCC recruitment onto endosomes and thereby release of WASH complex and tubules. (D) Release of regulated recycling tubules. TfR=transferrin receptor, C6-NBD-SM = C6-NBD-sphingomyelin.

In this study, we identified that COMMD10, as a subunit of the CCC complex, regulates endosomal recycling of ENaC. Our results also showed that extracellular Ca^2+^ leads to a vesicular F-actin accumulation while some other results showed that Ca^2+^ treatment increases COMMD10 protein level. Here, we propose that the increased endosomal actin-nucleation leads to stabilization of the CCC complex resulting in enhanced endogenous COMMD10 protein level. Both the increased endosomal actin-nucleation and COMMD10 protein level upregulates the cell surface activity of ENaC while an increase in endosomal actin-nucleation through the silenced CCC complex (in our case through COMMD10 knockdown) alone reduces the cell surface activity of ENaC. Taken together, COMMD10 protein level is required to be in a steady-state level for normal ENaC cell surface activity and thereby for normal body fluid volume and blood pressure.

## STAR★Methods

### Key Resources Table

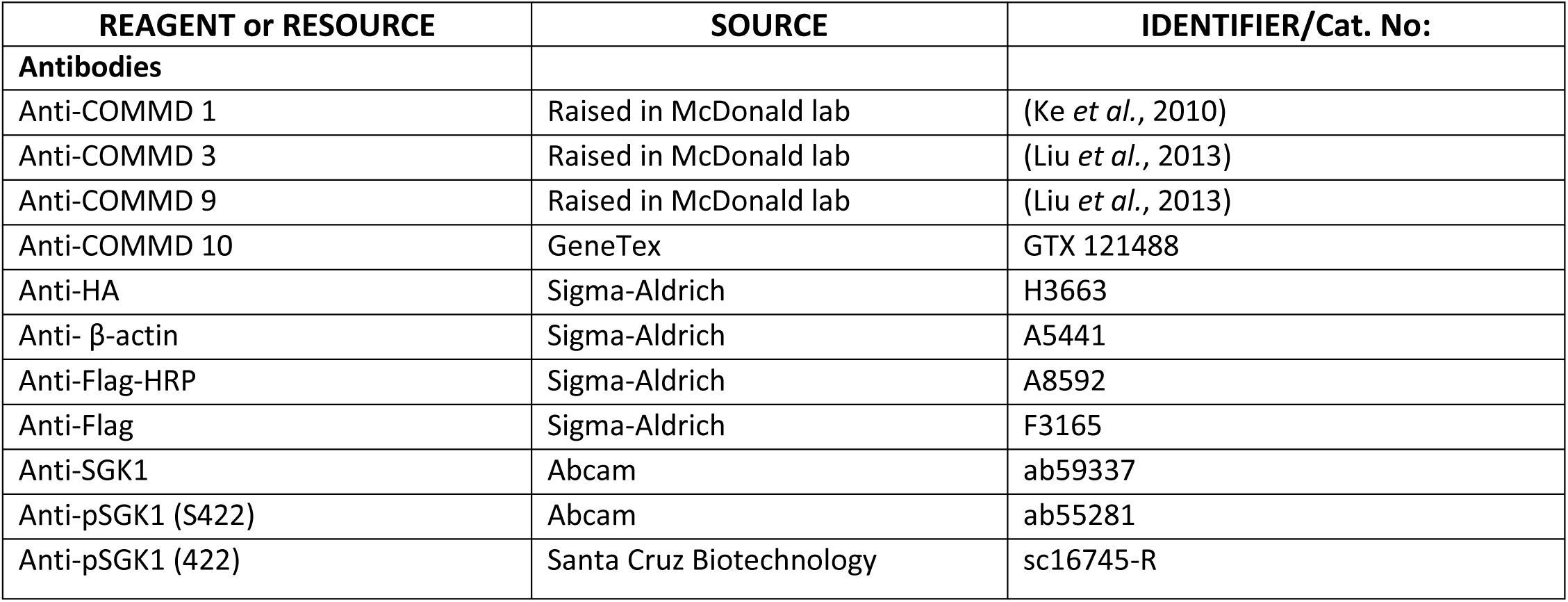
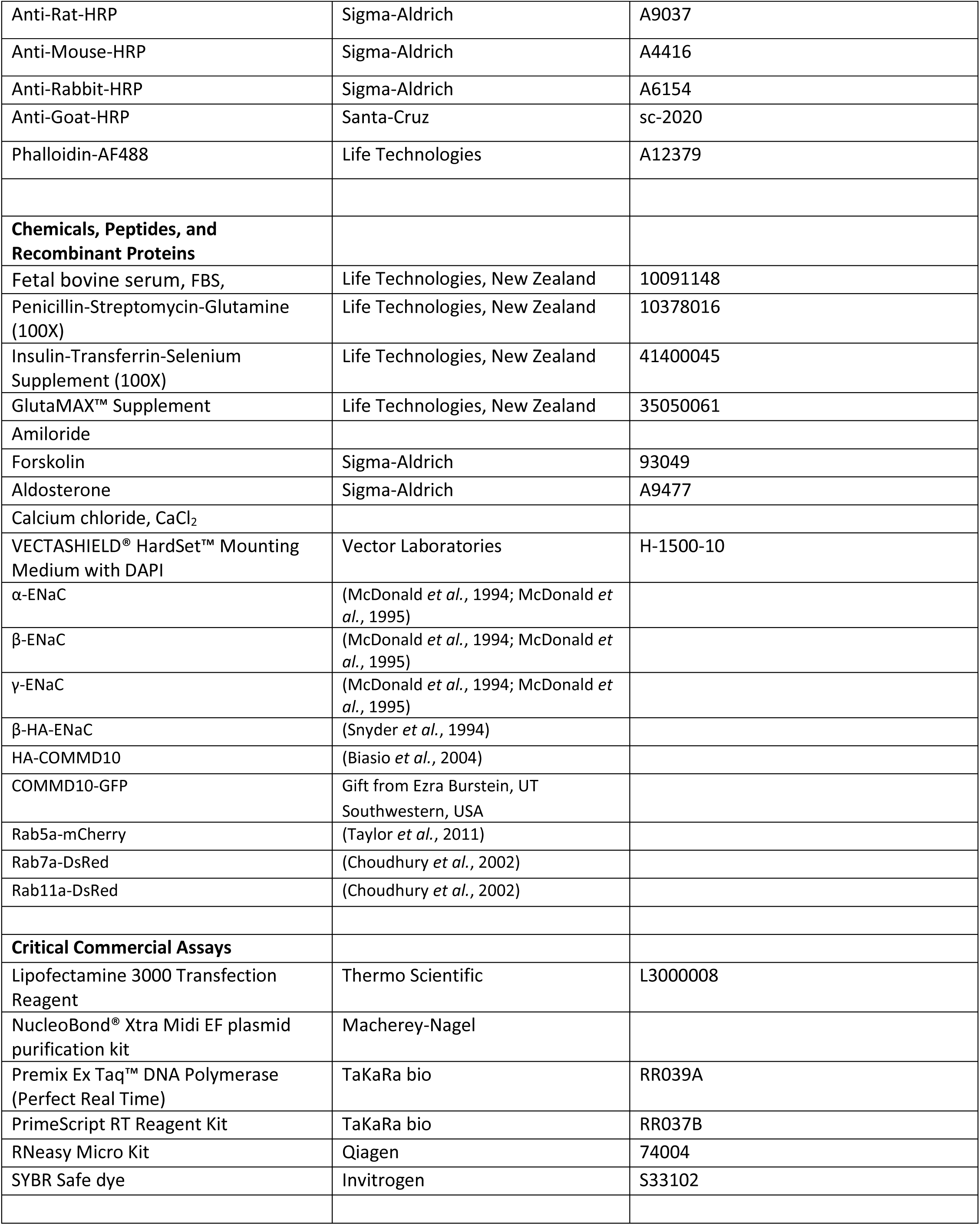
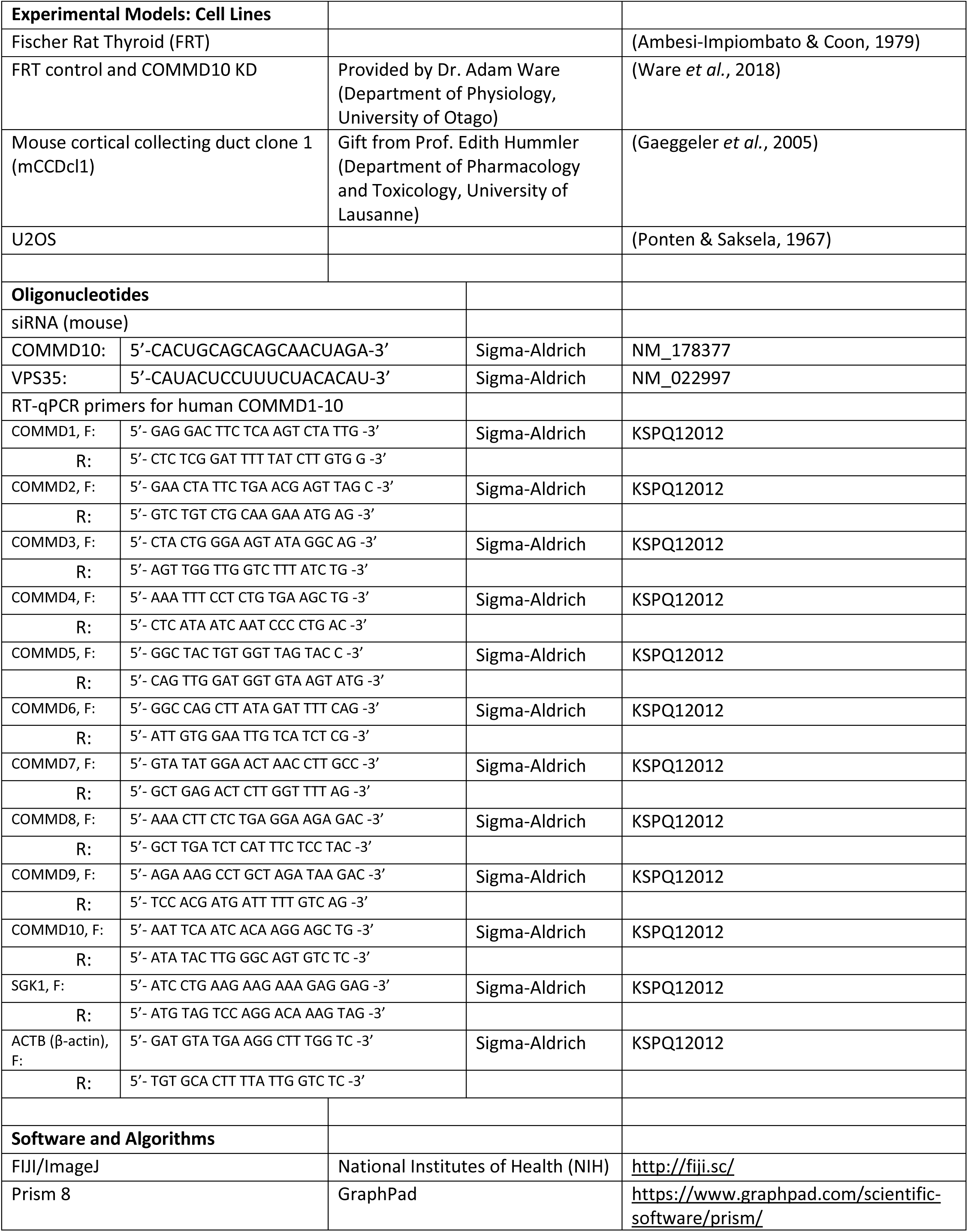
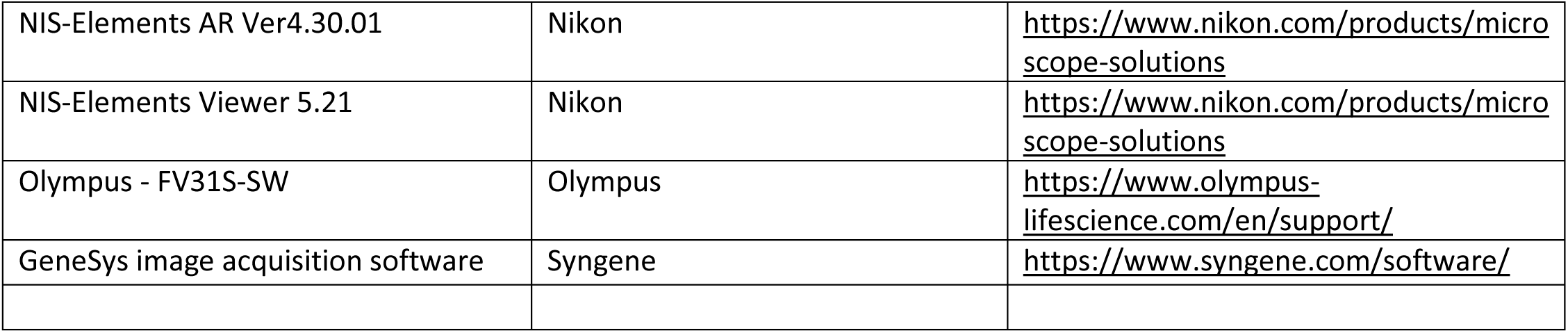

### Contact for Reagent and Resource Sharing

Further information and requests for resources and reagents should be directed to and will be fulfilled by the Lead Contact, Prof. Fiona McDonald.

### Method Details

#### Cell lines and plasmids

FRT cell lines (FRT control knockdown (KD) and COMMD10 KD) were maintained in Kaighn’s Modification F-12 media (Sigma Aldrich, New Zealand), supplemented with 10% (v/v) fetal bovine serum (FBS, Life Technologies, New Zealand), 1% (v/v) pen/strep (10 U/ml penicillin, and 100 μg/ml streptomycin (Life Technologies, NZ)). FRT control KD and COMMD10 KD cell lines were provided by Dr. Adam W. Ware (Ware *et al*., 2018). To preserve stable expression of short-hairpin RNA (shRNA) in FRT control and COMMD10 KD cell lines, the cells were treated with 2 µg/mL puromycin antibiotic at each passaging step. Silencing COMMD10 did not affect the cell growth or proliferation. For current measurements and for cell surface biotinylation assays, FRT cells were transfected with plasmids encoding α-,β-HA-,and γ-ENaC subunits. Mouse cortical collecting duct clone 1 (mCCDcl1) cells were provided by Prof. Edith Hummler (Department of Pharmacology and Toxicology, University of Lausanne). mCCDcl1 cells were cultured essentially as described previously (Gaeggeler *et al*., 2005; Mansley *et al*., 2015). U2OS cells were maintained in DMEM medium (Life Technologies, NZ) supplemented with 10% (v/v) FBS (Life Technologies, NZ), and 1% (v/v) penicillin–streptomycin (Life Technologies, NZ). For microscopy purposes, the FRT and U2OS cells were transfected with plasmids encoding COMMD10-GFP, Rab5a-mCherry, Rab7a- and Rab11-DsRed.

#### Western blot analysis

For direct Western blotting assays, T-TBS lysis buffer was used (1% Triton X-100 in Tris-Buffered Saline (TBS) (50 mM Tris·HCl and 150 mM NaCl, pH 7.4–7.5)) supplemented with protease inhibitors (10 μg/mL phenylmethylsulfonyl fluoride (PMSF), 2 μg/mL aprotinin, 2 μg/mL leupeptin and 1 μg/mL pepstatin) or 40 μl/mL 25x cOmplete™ Protease Inhibitor Cocktail (Sigma, Cat.No: 11836153001). After determining the protein concentrations, samples were boiled at 95°C for 5 minutes with addition of 5x Laemmli loading buffer (4% SDS, 20% glycerol, 0.004% bromophenol blue, 0.125M Tris-Cl, pH 6.8, 10% 2-mercaptoethanol). Depending on the protein mass, 8% - 15% SDS-PAGE gels were used to separate proteins. Protein samples and prestained protein marker (Precision Plus Protein™ Dual Color Standards, 1610374) were applied to the SDS-PAGE gel, run in a Hoefer Mini-Vertical system (Hoefer) at 100-150 V and transferred to polyvinylidene difluoride (PVDF) membrane by the semi-dry method. Membranes were incubated with 5% non-fat milk-TBST and blotted with primary antibody solution diluted in TBST for overnight at 4°C, then membranes were incubated with secondary antibody conjugated with horseradish peroxidase (HRP) for 1 hour at RT with 1:10,000 dilution. For chemiluminescent signal detection, Amersham™ ECL™ Prime Western Blotting Detection Reagent was used and blots were imaged by either exposing the membranes to X-ray film or in a Syngene Imager. Protein expression levels were normalized to β-actin or GAPDH expression.

#### Plasmid and siRNA transfection

SiRNA products were purchased from Sigma-Aldrich (Cat. No: NM_178377 or NM_022997). Using Lipofectamine^®^ 3000 (Thermo Scientific, Cat. No: L3000008), plasmids and/or siRNAs were transfected into FRT and mCCDcl1 cells following the manufacturer’s instructions. 24 h after seeding the cells, 1 μg DNA plus P3000 or 20 pmol siRNA without P3000 reagent was used to deliver into cells. Cells were incubated with the transfection mixture in FBS and antibiotic free medium for 5-6 hours which was followed by a change to fresh complete growth media. Cells were used for microscopy 48 h post transfection, for the endocytosis assay 24 h post transfection, for the recycling assay 120 h post transfection. Silencing COMMD10 or VPS35 did not affect the cell growth or proliferation.

#### Endocytosis assay

FRT cells stably expressing control shRNA and COMMD10 shRNA were transfected with plasmids encoding α-, β-HA, and γENaC subunits (1 µg of each) and after 48 hours, cells were incubated with cleavable biotin (EZ-link sulfo NHS-SS-biotin; Thermo Scientific Cat.No: 21331, final conc. 1 mg/mL) for 30 min after which the unreacted biotins were removed with quenching buffer (1% BSA in PBS) treatment for 1 min at 4°C. To allow surface proteins (including ENaC) to be internalized, cells were incubated for either 0, 2, 5, or 10 minutes at 37°C bathed with warm PBS++ (PBS supplemented with 1.5 mM MgCl_2_, 0.2 mM CaCl_2_, (pH 7.4)). After the incubation period, cells were treated with biotin stripping reagent L-glutathione (GSH) buffer (75 mM NaCl, 1 mM MgCl_2_, and 0.1 mM CaCl_2_, 50 mM GSH, 80 mM NaOH, and 10% FBS) six times for 15 min each to release the biotin groups from labelled, but uninternalized proteins during the incubation periods. Cells were lysed and after taking whole cell lysate (WCL) aliquots, the samples were incubated with NeutrAvidin beads to which the biotinylated and internalized proteins in the lysates were bound at 4°C overnight rotating. To pellet the bead-biotin-protein (including ENaC) complexes, lysates were centrifuged at 550 g for 5 min in a benchtop microcentrifuge. The pellets and also WCLs were analyzed by 10% SDS-PAGE gel and β-HA-ENaC detected using anti-HA antibody (Sigma-Aldrich, Cat.No. H3663, at a 1:2500 dilution).

#### RT-qPCR

For quantification of COMMD mRNA levels after aldosterone treatment, mRNA was extracted from mCCDcl1 cells lysing with TRIzol (Invitrogen) reagent and using a RNeasy Micro Kit (Qiagen, Cat No: 74004) according to the manufacturer’s guidelines. To erase any trace of genomic DNA (gDNA), an early (on column) DNase I treatment step was included in the protocol using the RNase-Free DNase Set supplied with the RNeasy Micro Kit. An ethanol precipitation method was used to extract mRNA for quantification of mRNA levels after calcium treatment. cDNA for real-time PCR was synthesized with 500 ng of extracted RNA using the PrimeScript RT Reagent Kit (Perfect Real Time) (TaKaRa bio, Cat. No: RR037B). RT-qPCR was performed with 10 ng cDNA in 10-μl reactions in triplicate or duplicate using Premix Ex Taq™ DNA Polymerase (TaKaRa bio, Cat. No: RR039A) and KiCqStart® SYBR® Green Primers (Sigma, Cat.No: KSPQ12012). The 2^−ΔΔCT^ method was used to analyze the relative changes in gene expression levels. Primers used in RT-qPCR are listed in the Key Resources Table.

#### Electrophysiological studies with FRT and mCCDcl1 epithelia

FRT cells were seeded on Snapwell filters (12 mm Snapwell, Corning® Costar, 3801) and grown in full media. 24 h after seeding, the cells were transfected with plasmids encoding α-, β-, and ϒ-ENaC (0.2 μg total). To measure amiloride-sensitive ENaC short circuit current (*Isc*-amiloride, or just *Isc*), the filters were mounted into an Ussing chamber 72 h after transfection. 1x Ringer’s solution (135 mM NaCl, 1.2 mM MgCl_2_, 4.2 mM K_2_HPO_4_, 0.6 mM KH_2_PO_4_, 10 mM HEPES, 1.2 mM CaCl_2_ – pH 7.4) was added to each side of the epithelium in the chamber. Two voltage sensing electrodes and two current passing electrodes were inserted into the ports (on each side of the epithelium). Chambers were maintained at 37°C and bubbled continuously with 100% O_2_ which fixed the pH at 7.4. The epithelia were clamped under short-circuit conditions and transepithelial resistance (Ω·cm^2^) was monitored by applying potential pulses (5 mV for 1 sec every 120 sec) across the epithelium. Upon current stabilization, 10 μM amiloride (final concentration) was added to the apical side of the epithelium. The *Isc* and transepithelial resistance were recorded by Acquire and Analyse 2.3 software (Physiologic Instruments).

Electrophysiological studies with mCCDcl1 cells was performed similar to FRT cells, but without transfection of ENaC subunits as mCCDcl1 epithelia endogenously express ENaC. Cells were grown for 6 days in growth medium which was changed to filter cup medium (without EGF and FBS) for the last 24 hours. The Snapwell inserts were mounted into an Ussing chamber and bathed with Ringer’s solution (pH 7.4) consisting of 120 mM NaCl, 25 mM NaHCO_3_, 10 mM D-glucose, 3.3 mM KH_2_PO_4_, 0.8 mM K_2_HPO_4_, 1.2 mM MgCl_2_, and 1.2 mM CaCl_2_. 10 μM amiloride was added to the apical bath to determine ENaC-mediated transepithelial short-circuit currents.

#### Recycling assay

An ENaC recycling assay was developed according to the assays conducted by Klemens et al. (Klemens *et al*., 2017). The method is based on measuring of *Isc* produced by ENaC that is delivered to the plasma membrane by cAMP stimulation. mCCDcl1 cells were transfected with control siRNA or siRNA against COMMD10 or VPS35 and after 120 hours, ENaC *Isc* was measured as described above. Upon the baseline current stabilization, 5 μM forskolin was added to the basolateral side of the epithelia and *Isc* continued to be measured. At the peak of FSK-stimulated *Isc* (which was achieved in 8-10 minutes and followed by a continuous but slight decline), 10 μM amiloride was added apically. The *Isc* value of the baseline *Isc* before FSK addition was subtracted from the value at the peak of FSK-stimulated *Isc*. *Isc* in control KD epithelia was normalized to one and *Isc* in COMMD10 or VPS35 KD was compared to the normalized control.

#### Aldosterone and calcium stimulation assay

For aldosterone stimulation experiments, mCCDcl1 cells were grown on permeable membrane filters (on 12 mm Snapwells for electrophysiological studies or on 24 mm Transwells for qPCR and Western blotting) were incubated with complete growth medium over a period of 5 days from seeding, to produce an epithelial monolayer with fully polarized epithelial features. On the fifth day after seeding, cells were incubated with filter cup medium, and subsequently, on the next day, cells were incubated for a further 24 h in FBS- and drug-free medium. Towards the end of the 24-hour incubation, the ‘stimulation medium’ was supplemented with 300 nM aldosterone (Sigma-Aldrich, Cat. No: A9477) in DMSO or just DMSO (as vehicle control) for various time periods.

For calcium stimulation experiments, mCCDcl1 or U2OS cells were treated with 0, 2, 4, or 6 mM CaCl_2_ (final concentrations of ∼2, ∼4, ∼6, or ∼8 mM CaCl_2_ in complete growth media, respectively) 24-hour after seeding the cells, and incubation continued for a total of 96 hours for Ussing assay, 72 hours for Western blotting and RT-qPCR, and 48 hours for microscopy.

#### Confocal imaging of fixed and live cells

To evaluate microfilament organization and intensity, F-actin was stained with phalloidin-AF488 (Life Technologies, A12379). FRT and U2OS control and COMMD10 KD cells were grown on 13 mm coverslips for 24 hours in complete media. U2OS cells for calcium treatment, were further incubated for 48 hours with 2 mM CaCl_2_ treatment. Cells were fixed in 3.7% formaldehyde solution in PBS for 10 minutes at RT and permeabilized with 0.1% Triton X-100 in PBS for 3-5 min. After blocking with PBS + 1% BSA for 20–30 min, microfilaments of the cells were stained with 0.17 µM of phalloidin-AF488 (final concentration) following the manufacturer’s instructions. After washing in PBS, the samples were mounted with VECTASHIELD^®^ HardSet™ Mounting Medium with DAPI (Vector Laboratories, H-1500).

To visualize COMMD10 protein on Rab5-, Rab7-, and Rab11-positive endosomes in living cells, Rab5a-mCherry, Rab7-DsRed, or Rab11-DsRed together with COMMD10-GFP plasmids were coexpressed in U2OS and FRT COMMD10 stable knockdown cells. To avoid overexpression-related COMMD10 aggregation, U2OS and FRT COMMD10 KD cells were used. Images of living cells were acquired after less than 24 hours using a Nikon A1R inverted confocal microscope equipped with 60× PlanApo (1.49 NA, oil immersion) objective or an Olympus FV3000 microscope equipped with a temperature-, humidity-, and CO_2_-controlled chamber and 60× objectives. Analyses were performed on endosomes with extended tubules observed mostly in time lapse movies to ensure the tubule-like structures were attached to endosomes rather than being distinct structures moving or passing around them. Images were analyzed using FiJi software. For estimating COMMD10 enrichment, a line was drawn along the endosome and tubule-like structure, and the Plot Profile function was used to measure pixel values. For obtaining the average value plot across multiple endosome-and-tubule-like structures, the length of these compartments was firstly standardized and then the linear pixels were normalized to the maximum fluorescence value of each compartment and averaged in GraphPad Prism (v.8.3.1).

#### Quantification and Statistical Analysis

All statistical analysis was performed using GraphPad Prism v8.3.1. For most analyses, one sample t-test for normalized data, unpaired t-test for unnormalized pair of data, and two-way ANOVA for a variable measured in three or more groups was used. The P values of the statistical tests were regarded as significant if P<0.05, P<0.01, P<0.001, or P<0.0001 and indicated as *, **, ***, or ****, respectively. Data were presented as mean ± standard deviation (SD).

For means of pixel distributions and average *Isc* traces, standard error of the mean (SEM) was used. Individual experiment numbers are indicated with “N” while biological replicates were indicated with “n”.

## Acknowledgments

This study was funded by the University of Otago Doctoral Scholarship to S.R.R. and was supported by Department of Physiology and University of Otago Division of Health Sciences Travel Grant. We gratefully acknowledge Prof. Edith Hummler, University of Lausanne for mCCDcl1 cells and Dr. Adam Ware for COMMD10 stable knockdown cells.

## Author Contributions

S.R.R. and F.J.M. conceived and designed the study and wrote the manuscript. S.R.R. performed most of the experiments and data analyses.

## Supplemental Information

