## Supplemental Information for "COMMD10 Regulates Endosomal Recycling of Epithelial Sodium Channel (ENaC)"

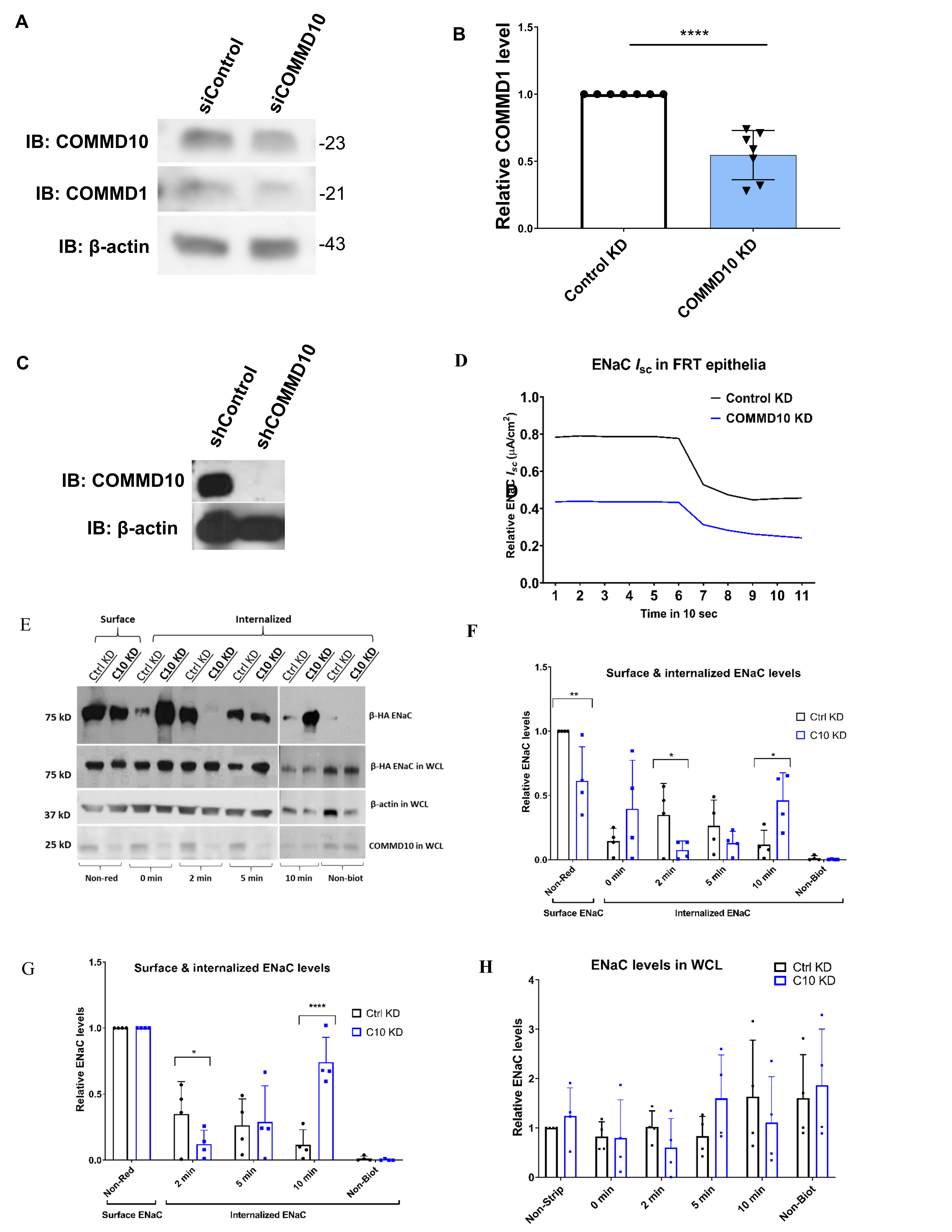

Figure S1. COMMD10 KD destabilizes CCC complex and thereby impairs ENaC is accumulated intracellularly. Related to Figure 1.

(A) Representative blots for COMMD1 protein together with COMMD10 and loading control β-actin. mCCDcl1 cells were transfected with control or COMMD10 siRNA.

(B) Pooled relative protein levels of COMMD1 that was decreased in COMMD10 KD cells. Student’s t test. **P˂0.01, ****P˂0.0001, mean ± SD. N=3. n=7.

(C) Representative western blot for COMMD10. FRT cells stably expressing control and COMMD10 short hairpin RNA (shControl and shCOMMD10) were lysed and analyzed on a 15% SDS-PAGE gel. N=3, n=7

(D) Representative current tracings for *Isc* in FRT control and COMMD10 KD (C10 KD) epithelia. N = 5

(E-F) FRT control and COMMD10 knockdown cells were transfected with plasmids encoding α-,β-HA, and γ-ENaC subunits. Cell surface proteins including ENaC were biotinylated with cleavable biotins that followed by incubation for 0, 2, 5, and 10 minutes at 37°C to allow internalization of cell surface proteins. After the incubation step, non-internalized biotins on the cell surface were reduced with L-glutathione and then biotin-bound proteins including ENaC was precipitated with NeutrAvidin beads and analyzed by Western blotting. (E) Representative western blots β-HA ENaC, COMMD10 and β-actin. The upper panel shows blots of cell surface and internalized β-HA ENaC while the lower rows of blots are from whole cell lysates (WCL). *In two of the experiments, biotins were found to bind intracellular proteins of 0 min COMMD10 KD cells, possibly due to cell membrane damage. (F) Relative endocytosis levels. β-HA ENaC cell surface level in non-reduced control cells was normalized to 1. Then cell surface level of ENaC in COMMD10 KD cells and internalized ENaC values of each time point were normalized to that of non-reduced control cells. Two-way ANOVA (applying Fisher's LSD test) were used to compare and demonstrate any difference between control and COMMD10 KD surface and internalization at each time point. Internalized ENaC in COMMD10 KD cells was significantly accumulated intracellularly with 10 mins of incubation.

(G) Internalized ENaC levels proportional to the cell surface ENaC levels, where the surface ENaC levels of both of non-reduced control and COMMD10 KD cells were normalized to 1 and the internalized ENaC levels of each time point were compared to the normalized control or COMMD10 KD surface levels, respectively.

(H) Whole cell lysate ENaC levels that showed no significant changes in COMMD10 KD cells compared to that of control KD cells as expected.

*Non-Red*—non-reduced, *Non-Biot* —non-biotinylated. Data shown as mean ± SD relative to normalized ENaC surface level of control KD epithelia. Two-way ANOVA (Mult. comp.-Fisher's LSD test). N=4. *P˂0.05, **P˂0.01.

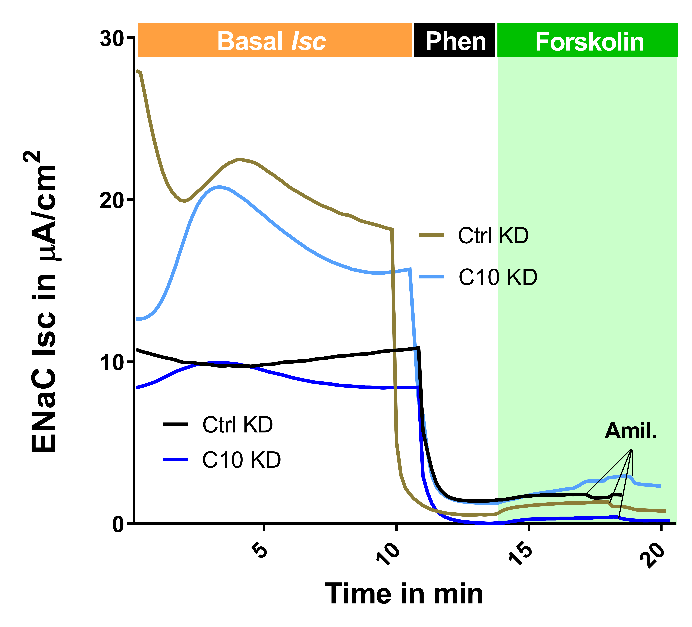

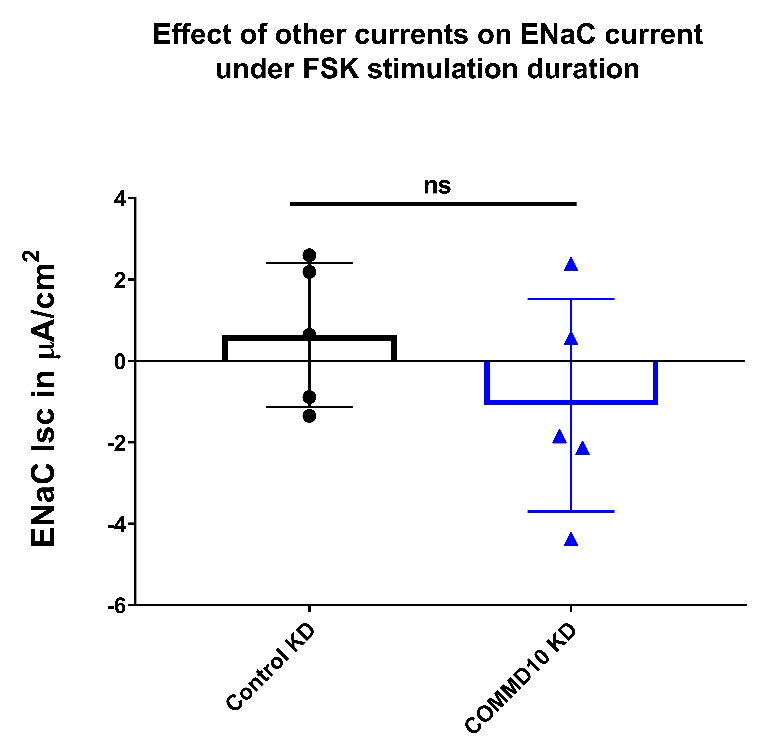

**B**

**A**

**Figure S2. COMMD10 KD does not alter other currents during FSK stimulation. Related to Figure 1.**

(A) Upon stabilization of basal ENaC *Isc*, 25 μM phenamil was added apically which was followed by 5 μM FSK addition basolaterally. Forskolin slightly increased the current in both the control and COMMD10 KD epithelia, however, upon addition of amiloride 5 min after forskolin addition, most of the FSK-induced current was reduced suggesting that most of the FSK-stimulated current is produced by ENaC newly inserted into apical membrane but not blocked yet by phenamil. N=1, n=2.

(B) Pooled results from the analysis of ENaC *Isc* at 3 min after adding amiloride at the peak of the current (the last series of traces in Figure 1C) confirming that the reduction in other currents in COMMD10 KD is not significant (0.64 ± 1.8 for control KD vs -1.1 ± 2.6 μA/cm^2^ for COMMD10 KD epithelia, P>0.05, N=3, n=5).

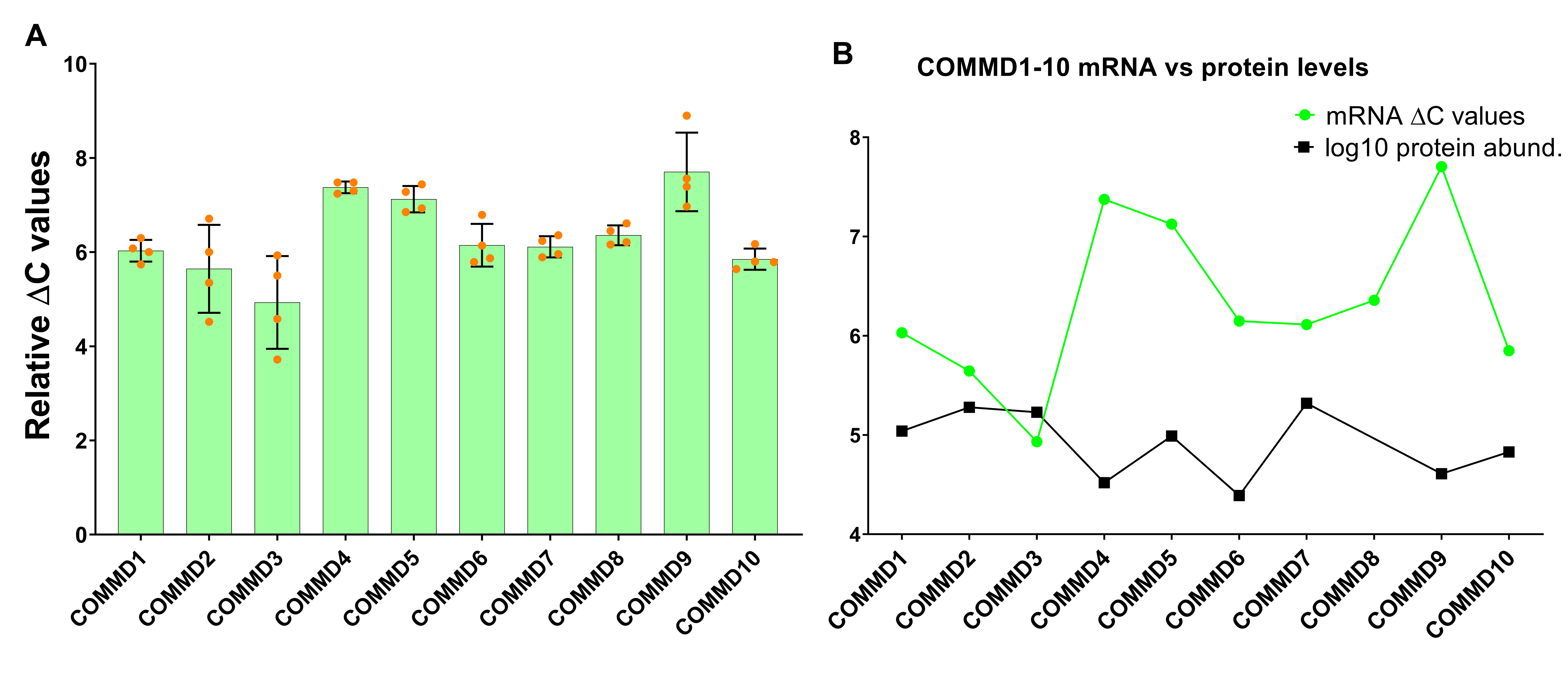

Figure S3. COMMD1 is reduced in COMMD10 attenuated mCCDcl1 cells. Related to Figure 4.

(A) mRNA levels of COMMD1-10 (given in ΔC value – Ct value of a COMMD minus Ct value of β-actin of the same sample) in mCCDcl1 epithelia. The results show uneven levels of COMMD1-10 mRNA levels. mean ± SD. N=4.

(B) Comparison of mRNA ΔC values in mCCDcl1 epithelia with logarithmic (10) protein abundance of COMMD1-10 in mpkCCD epithelia (Yang *et al.*, 2015).

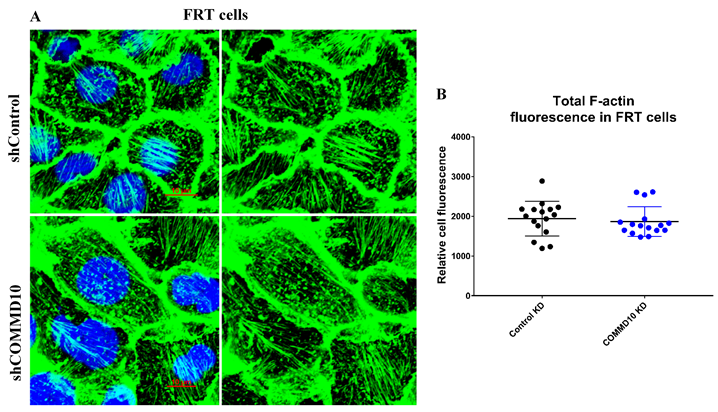

**Figure S4. COMMD10 KD doesn’t change microfilament organization of FRT cells. Related to Figure 6.**

(A) Representative images of stained F-actin in FRT control and COMMD10 KD cells. Scale bar = 10 μM.

(B) Quantitative analysis of fluorescence intensity of actin filaments showing actin organization and intensity in control and COMMD10 KD FRT cells. Student’s *t*-test, P=0.62, mean ± SD. n=272 for control KD cells, n=253 for COMMD10 KD cells.
